# Peroxide antimalarial drugs target redox homeostasis in *Plasmodium falciparum* infected red blood cells

**DOI:** 10.1101/2021.09.10.459878

**Authors:** Ghizal Siddiqui, Carlo Giannangelo, Amanda De Paoli, Anna Katharina Schuh, Kim C. Heimsch, Dovile Anderson, Timothy G. Brown, Christopher A. MacRaild, Jianbo Wu, Xiaofang Wang, Yuxiang Dong, Jonathan L. Vennerstrom, Katja Becker, Darren J Creek

## Abstract

*Plasmodium falciparum* causes the most lethal form of malaria. Peroxide antimalarials based on artemisinin underpin the frontline treatments for malaria, but artemisinin resistance is rapidly spreading. Synthetic peroxide antimalarials, known as ozonides, are in clinical development and offer a potential alternative. Here, we used chemoproteomics to investigate the protein alkylation targets of artemisinin and ozonide probes, including an analogue of the ozonide clinical candidate, artefenomel. We greatly expanded the list of protein targets for peroxide antimalarials and identified significant enrichment of redox-related proteins for both artemisinins and ozonides. Disrupted redox homeostasis was confirmed by dynamic live imaging of the glutathione redox potential using a genetically encoded redox-sensitive fluorescence-based biosensor. Targeted LC-MS-based thiol metabolomics also confirmed changes in cellular thiol levels. This work shows that peroxide antimalarials disproportionately alkylate proteins involved in redox homeostasis and that disrupted redox processes are involved in the mechanism of action of these important antimalarials.

**Importance:** The frontline treatments for malaria are combination therapies based on the peroxide antimalarial, artemisinin. Concerningly, artemisinin resistance has emerged in malaria-endemic regions, and now poses a major threat to malaria treatment and eradication efforts. New medicines are urgently required to replace the artemisinins, and some of the most advanced candidates are the fully synthetic peroxide antimalarials, OZ277 (arterolane) and OZ439 (artefenomel). The mechanism of action of peroxide antimalarials involves the reductive activation of the peroxide bond by intra-parasitic haem, but there is no consensus regarding the specific protein targets of the resulting radical species for artemisinins and/or the ozonides. This study provides a comprehensive and unbiased chemoproteomic profile of over 400 target proteins, and confirms the specific impact of peroxide antimalarials on redox metabolism. The key role of redox targets is particularly relevant considering that the mechanism of artemisinin resistance appears to involve modulation of peroxide activation and redox homeostasis.

## Introduction

Malaria is a major global health challenge and an estimated 409 000 deaths and 229 million new malaria cases occurred worldwide in 2019^1^. The majority of malaria-related mortality is due to infection with the deadliest parasite species, *Plasmodium falciparum*. Currently, antimalarial drugs are one of the most powerful tools for combating malaria. The first-line treatment options for uncomplicated malaria are artemisinin-based combination therapies (ACTs). However, the emergence and spread of parasite strains that are resistant to multiple antimalarial drugs is a major challenge to malaria treatment.

The clinically-used artemisinins, such as artemether, artesunate and dihydroartemisinin (DHA), are semisynthetic derivatives of the natural product artemisinin and possess a 1,2,4-trioxane core incorporating a peroxide bond that is essential for activity^2^. Artemisinins clear *P. falciparum* infections rapidly and provide prompt resolution of malaria symptoms in patients with both uncomplicated and severe infections^3^. However, artemisinins are limited by poor biopharmaceutical properties and short *in vivo* half-lives (typically < 1 h in humans)^4^. This necessitates a 3-day treatment regimen and coadministration with a long acting partner drug.

To overcome some of these limitations, the peroxide bond of artemisinin was used as inspiration for the design of synthetic peroxide antimalarials, collectively known as ozonides. The first-generation ozonide, OZ277 (arterolane), was found to be rapidly acting and exhibits similar blood-stage activity to clinically-used artemisinin derivatives, both *in vitro* and *in vivo*^5^. In 2012, OZ277 was the first synthetic peroxide antimalarial to be approved for clinical use. Continued development of the ozonides led to a second-generation series of compounds that had a significantly improved *in vivo* exposure profile and maintained the potent *in vitro* and *in vivo* activity of the first-generation compounds^6–7^. The second generation ozonide clinical candidate, OZ439 (artefenomel), is the first long half-life peroxide antimalarial to be tested clinically^8–9^ and has reached advanced stages of development in combination with ferroquine.

Artemisinins and ozonides are thought to act through similar mechanisms as both classes require the peroxide pharmacophore for activity^2, 10–11^. The proposed mechanism of action involves initial bioactivation by haem released through parasite digestion of haemoglobin^12–15^. This process results in cleavage of the peroxide bond and generation of carbon-centered radicals ^11, 16^ that alkylate haem^17–18^ and proteins^19–26^ and induce widespread oxidative stress^27–31^. Death of the parasite likely results from disruption to multiple vital processes, including haemoglobin degradation in the food vacuole^32^.

Concerningly, parasites with decreased sensitivity to artemisinins have emerged in the Greater Mekong Subregion^33^, and more recently in eastern India^34^, Africa^35^ and Papua New Guinea^36^. Clinically this manifests as delayed parasite clearance following treatment with an ACT^37–38^ and can lead to approximately 50% treatment failure in areas with concomitant partner drug resistance^39–40^.

Delayed parasite clearance is associated with point mutations in the propeller domain of the *P. falciparum* Kelch 13 protein^41^ and decreased parasite susceptibility to short exposures of artemisinin *in vitro*^42–43^. Although the delayed parasite clearance phenotype does not represent complete resistance^44^, the emergence of these parasites poses a major threat to global malaria control efforts. For this reason, it is critically important to understand the mechanism of action of antimalarial peroxides, with a view to overcoming resistance. The potential for cross-resistance between artemisinins and ozonides has also been the subject of considerable debate^45^ and some reports suggest that the second-generation series of ozonides, e.g. OZ439, may be less susceptible to the mechanisms underpinning decreased artemisinin sensitivity^46–48^. This raises the prospect that first generation ozonides, second generation ozonides and artemisinins could alkylate unique targets within the parasite. Previous click chemistry-based proteomics studies aimed at identifying the protein targets of peroxides have shown that artemisinin and simple ozonide probe compounds (containing the 1,2,4-trioxolane core flanked by a cyclohexane moiety on one side and an adamantane group on the other) share a common alkylation signature^22^. Whereas, others have proposed that protein alkylation is indiscriminate and stochastic^20, 23^.

Here, we used chemical proteomics to directly compare the temporal protein alkylation profile of clickable peroxide probes based on artemisinin, the simple ozonide core structure and the ozonide clinical candidate, OZ439. We employed a unique data independent acquisition mass spectrometry (DIA-MS) approach and extensive controls to generate an extended and robust list of protein alkylation targets for peroxide antimalarials, identifying redox homeostasis proteins as key targets.

The impact to redox homeostasis was confirmed using a ratiometric redox sensor consisting of human glutaredoxin 1 fused to a reduction-oxidation sensitive GFP (hGrx-1-roGFP2)^49^ and targeted LCMS-based thiol metabolomics. Taken together, we demonstrate that the mechanism of action of peroxide antimalarials involves disproportionate alkylation of proteins involved with redox processes and significant disruption to *P. falciparum* redox homeostasis.

## Results

### Design and synthesis of alkyne peroxide probes for protein target identification

To profile and identify the protein targets of peroxide antimalarials, we designed and synthesised a series of ozonide and artemisinin probe compounds with a clickable alkyne tag (Figure 1A). The alkyne tag can be further appended with agarose beads through click chemistry, allowing the targets of the peroxide probes to be affinity purified for mass spectrometric analysis (Figure 1B). The active peroxide probes (AA2, OZ727 and OZ747) containing alkyne functionality retained potency against *P. falciparum* 3D7 parasites as determined by their IC_50_ *in vitro*, whereas the alkyne-modified non-peroxide control probe, carbaOZ727, was inactive (Figure 1A). These findings confirmed that addition of the clickable alkyne tag does not interfere with probe efficacy and that antimalarial activity is peroxide-bond dependent.

**Figure 1.**
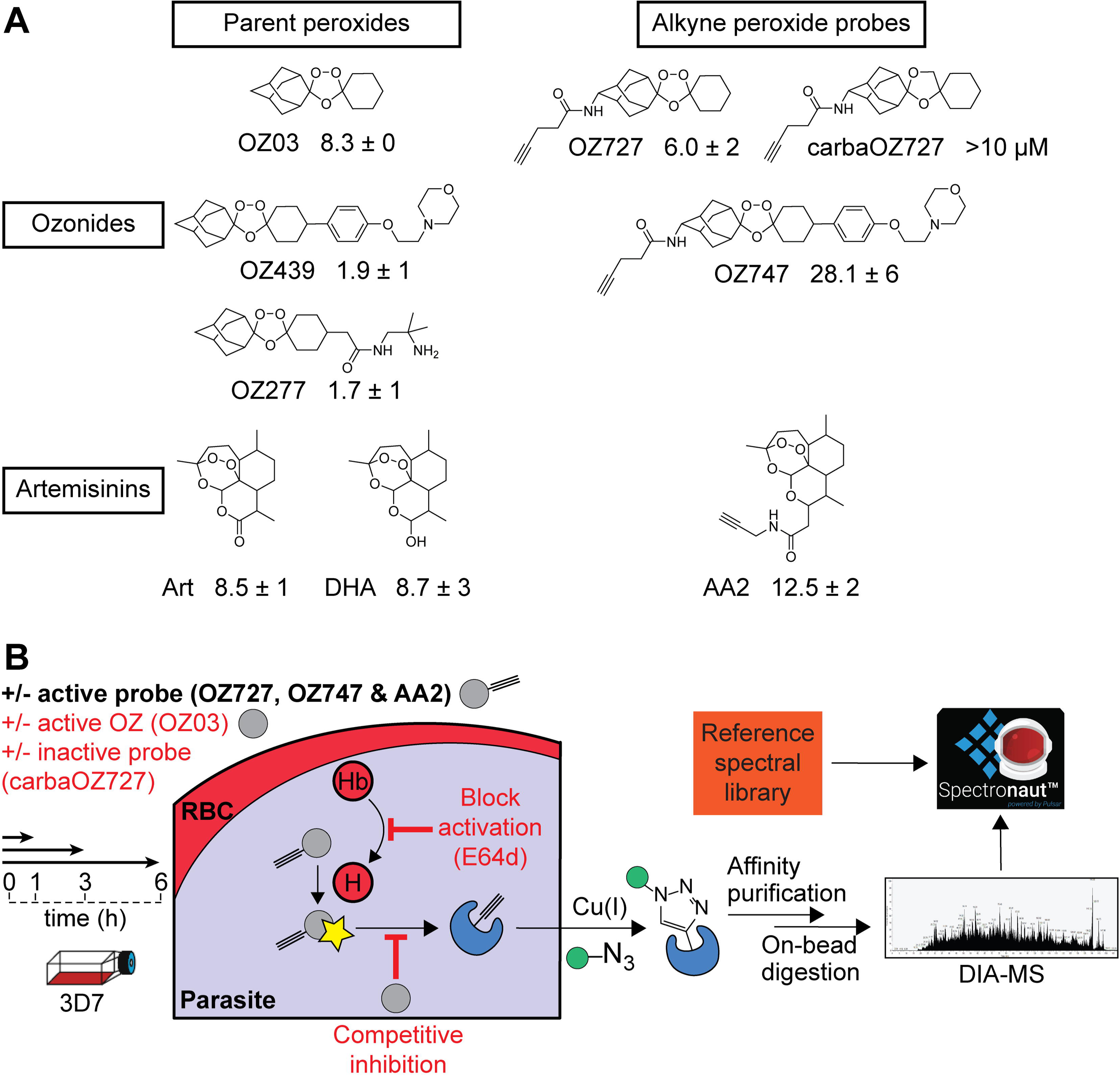
Identification of peroxide antimalarial molecular targets by click chemistry-based chemical proteomics. (A) Molecular structures of clickable alkyne probes OZ727, OZ747, AA2 and carbaOZ727, and their associated non-clickable controls OZ03, OZ277, OZ439, Art (artemisinin) and DHA (dihydroartemisinin). Activity of test compounds against *P. falciparum* 3D7 (nM) is shown as the mean ± SEM of data from three independent replicates with at least two technical repeats. (B) General workflow of the copper-catalysed click chemistry approach used to identify protein targets of peroxide antimalarials. *P. falciparum* parasites are incubated with the alkyne probes, which get activated (star) by free haem (H) derived from haemoglobin (Hb) digestion to form free radicals that can alkylate protein targets in the cell. The alkyne-modified proteins are affinity purified by covalent attachment to azide agarose beads using copper-catalysed click chemistry, and identified using data independent acquisition (DIA) LCMS/MS. The complex spectra obtained from the MS is then processed using a reference in-house *P. falciparum* spectral library in Spectronaut^TM^ Software. As an additional strategy to identify biologically relevant protein targets, peroxide activation was blocked *in situ* with the falcipain haemoglobinase inhibitor, E64d, which antagonises the antimalarial effects of peroxides, and the non-clickable parent ozonide, OZ03, served as a competitive inhibitor of probe binding *in situ* to distinguish specific versus non-specific protein hits. Control conditions are shown in red text.

We employed a time course treatment and comprehensive control strategy for our affinity purification experiments to ensure specific and high confidence identification of biologically relevant protein targets (Figure 1B). The inactive (clickable) non-peroxide probe, carbaOZ727, and non-clickable parent ozonide, OZ03, acted as controls for false positive protein identifications. OZ03 also served as a competitive inhibitor of probe binding *in situ* to distinguish specific versus non-specific protein hits (Figure 1B). As parasite degradation of haemoglobin provides the iron source for peroxide activation, we blocked peroxide activation *in situ* with E64d (Figure 1B), a falcipain haemoglobinase inhibitor known to antagonise the antimalarial effects of peroxides^12–13, 27^, to further eliminate proteins unlikely to be involved with the action of peroxides in parasite killing.

### Identification of protein targets of peroxide antimalarials

For the identification of peroxide covalent binding targets, synchronised trophozoite cultures were incubated for 1-6 h with 300 nM of the alkyne probes, OZ727, OZ747 or AA2. Following parasite isolation and protein extraction, alkyne labelled proteins were affinity purified by copper catalysed click chemistry onto azide agarose beads and protein targets were identified with DIA-MS (Figure 1B). Protein intensities were used to provide semi quantitative analysis of protein abundances for a total of 928 parasite proteins identified across all sample groups (Supplementary Data 1). Proteins in the OZ727, OZ747 and AA2 groups with a fold-change ≥ 2 compared to the DMSO, carbaOZ727 and time-matched OZ03 controls, and with a p-value < 0.05 (Mann-Whitney *U* test) compared to carbaOZ727 or DMSO, were considered as protein targets. After removal of false positive protein identifications and non-specific protein hits, a total of 182, 94 and 261 parasite proteins were respectively identified as targets of OZ727, OZ747 and AA2 after 1 h of treatment (at least four independent experiments) (Figure 2A). OZ727 labelling was markedly decreased by co-incubation with increasing concentrations of OZ03 (active non-clickable parent ozonide), indicating that the engineered alkyne probes alkylate the same targets as their non-clickable parents (Figure 2A and Supplementary Data 1). Furthermore, pre-incubation of parasite cultures with E64d drastically decreased labelling of proteins with OZ727 (Figure 2A and Supplementary Data 1), consistent with reports that inhibiting haemoglobin digestion abrogates peroxide activity^12–13, 27, 50^. Taken together, these results indicate the alkyne probes are pharmacologically similar to the parent compounds and the protein targets identified in this study are associated with their antimalarial peroxide activity^51^.

**Figure 2.**
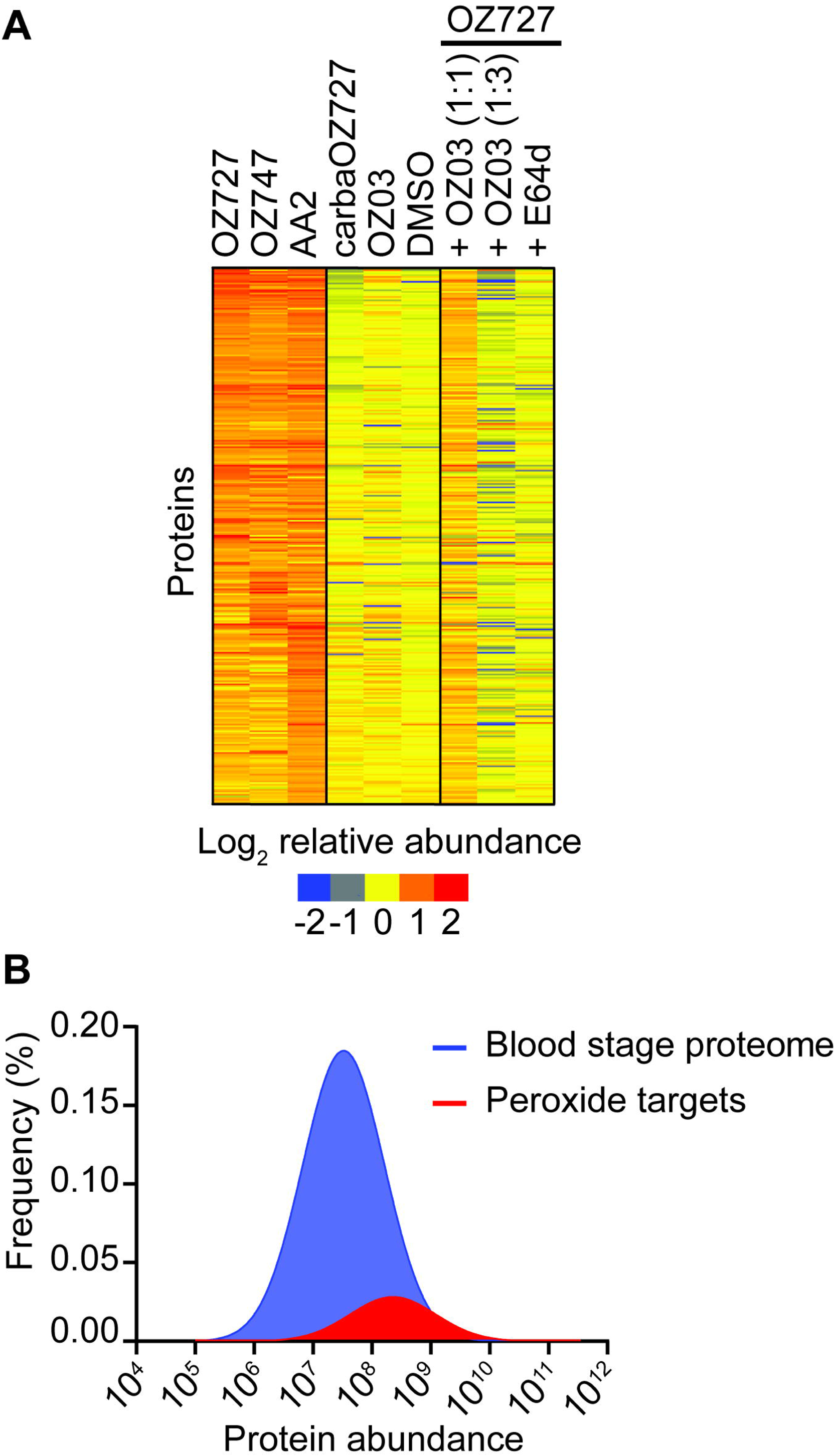
Protein binding targets of peroxide antimalarials in *P. falciparum*. (A) Heatmap representation of all protein binding targets of OZ727, OZ747 and AA2 following treatment of *P. falciparum* infected RBCs for 1 h (all 300 nM). Heatmap analysis was performed by plotting the average log transformed protein abundance for each protein and normalising by the mean for that protein across all samples. Proteins in the OZ727 (n = 6), OZ747 (n = 5) and AA2 (n = 4) groups with a fold-change ≥ 2 compared to the DMSO (n = 9), carbaOZ727 (n = 7) and OZ03 (n = 3) controls, and with a p-value < 0.05 (Mann-Whitney *U* test) compared to carbaOZ727 or DMSO, were considered as protein binding targets. Protein labelling by OZ727 was decreased in the presence of increasing concentrations of the active non-clickable parent ozonide, OZ03 (n = 3-4), and by pre-incubation of parasite cultures with the cysteine haemoglobinase inhibitor, E64d (n = 4). For additional protein information see Supplementary Data 1. (B) Histogram showing that the 436 proteins identified as peroxide binding targets across all study timepoints (red) are distributed across the *P. falciparum* blood stage protein abundance range and not limited to the most abundant proteins within the parasite. The frequency distribution of proteins in the *P. falciparum* blood stage proteome (Siddiqui et al., unpublished data) is shown in blue.

Importantly, the proteins labelled by OZ727, OZ747 and AA2 were not simply the proteins that are reproducibly detected as the most abundant proteins in untargeted proteomic studies of *P. falciparum* parasites (Figure 2B). Approximately 45% of the proteins identified as targets of peroxides were among the top 500 most abundant proteins in the *P. falciparum* blood stage proteome. This contrasts with previous peroxide chemical proteomics studies, where between 60% and 80% of proteins identified were among the most abundant proteins^21, 23^. A significant proportion of the proteins identified as peroxide targets were also detected by Wang *et al*.^23^ (55%) and Ismail *et al*.^21^ (36%) (Figure S1A). However, only 13 proteins were common between all three studies and these were all highly expressed *P. falciparum* blood stage proteins, with the exception of the putative Fe-S cluster assembly protein DRE2 (PF3D7_0824600) (Figure S1B). In another peroxide chemical proteomic study, where the NF54 parasite line was used, 25 protein targets were identified with 100 ng/mL of the chemical probes (similar to the probe concentration used in this study). Of the 25 *P. falciparum* proteins identified as targets, 8 were also identified in our list of targets; however, these 8 proteins were not the same as the common 13 proteins identified across the other studies ^20^. In addition to these commonly identified proteins, our unique data acquisition approach with extensive controls added a further 352 biologically relevant proteins to the list of peroxide targets in *P. falciparum* (Figure S1B).

In our dataset, approximately 30% of the combined 435 proteins alkylated by OZ727, OZ747 and AA2 were identified as targets for all three compounds. A range of proteins were also identified as unique targets for OZ727 (79 proteins), OZ747 (35 proteins) or AA2 (59 proteins). For example, AA2 alkylated several proteins involved in vesicle-mediated transport (PF3D7_0303000, PF3D7_1231100 and PF3D7_1429800) that were not binding targets for OZ727 or OZ747. Similarly, several proteins involved in nucleotide biosynthesis (PF3D7_0206700, PF3D7_0629100, PF3D7_1012600 and PF3D7_1437200), an essential pathway for parasite growth and survival, were only alkylated by OZ727, while OZ747 uniquely alkylated proteins involved in ion transport (PF3D7_0715900 and PF3D7_1362300). Some of the major proteins previously identified as targets of artemisinin and simple ozonide probe compounds, such as translationally controlled tumour protein (PF3D7_0511000)^52^, actin 1 (PF3D7_1246200), ornithine aminotransferase (PF3D7_0608800) and the glycolytic enzymes, lactate dehydrogenase (PF3D7_1324900), enolase (PF3D7_1015900) and glyceraldehyde-3-phosphate dehydrogenase (PF3D7_1462800)^21–23^, were confirmed as targets for AA2 and OZ727, but were not alkylated by the OZ439 analogue, OZ747.

We performed gene ontology (GO) enrichment analysis to identify the major cellular processes and compartments targeted by the peroxide antimalarials. These data showed that the targets of OZ727, OZ747 and AA2 were significantly enriched in several cellular components, including the cytosol, ribosome, nucleus and food vacuole (Supplementary Data 2). Notably, the food vacuole (GO:0020020) was the most significantly enriched compartment for both OZ727 (p-value = 9.90E-12) and AA2 (p-value = 1.30E-15) at the 1 h timepoint and was the second most significantly enriched compartment for OZ747 (p-value = 0.00019) (Supplementary Data 2). Pre-treatment with E64d and co-incubation with increasing concentrations of OZ03 inhibited OZ727 binding to food vacuole proteins (Figure S2), confirming that these targets are important for peroxide activity. These findings are consistent with peroxides initially acting by disrupting parasite haemoglobin digestion^32^, an essential process that occurs within the parasite food vacuole. GO analysis further revealed that peroxide protein targets were involved in numerous essential biological processes of the parasite (Figure 3 and Supplementary Data 3). Several significantly enriched biological processes were identified in redox homeostasis, metabolism, nuclear transport and stress response (Figure 3). To control for non-specific protein binding, enrichment analysis of proteins identified in the control samples found GO terms associated with nuclear transport and stress response, suggesting that those functions may not be specifically targeted by these drugs. However, enrichment of GO terms associated with redox homeostasis and metabolism were specific to the peroxide-treated parasites and not the controls (Figure 3 and Supplementary Data 3). Of these, cell redox homeostasis (GO:0045454) was one of the most significantly enriched biological process GO terms for both OZ727 (p = 0.00015) and AA2 (p = 0.0016) after 1 h and was not enriched in the control dataset (Figure S3 and Supplementary Data 3).

**Figure 3.**
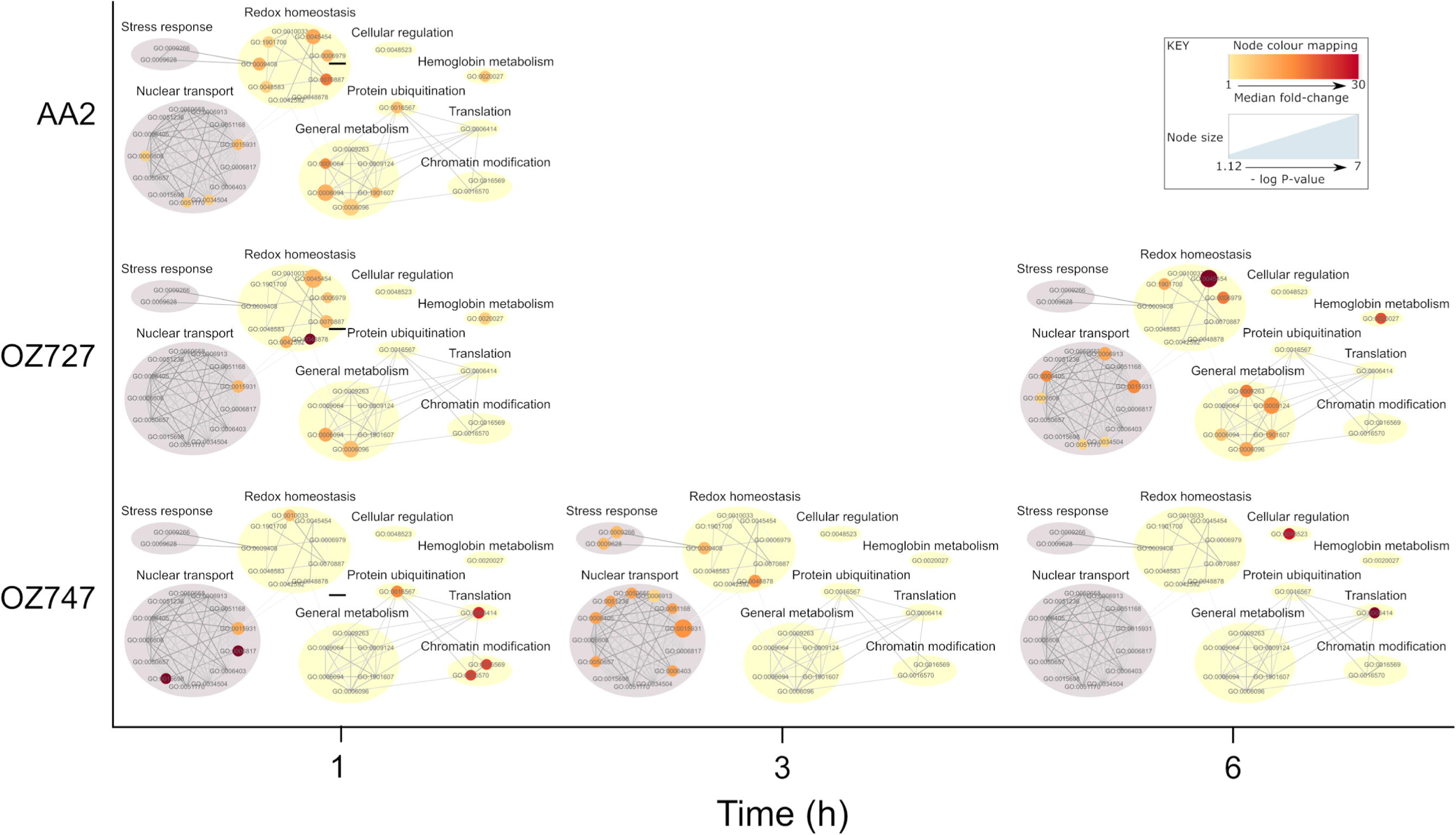
Gene ontology (GO) biological process enrichment of *P. falciparum* proteins covalently interacting with peroxide antimalarials. Peroxide protein targets are involved in multiple vital biological processes. GO analysis was conducted using topGO. PlasmoDB GO terms were used for GO term mapping and the elim algorithm was applied to navigate the topology of the GO graph. The Fisher exact test was used for testing enrichment of GO terms. *P. falciparum* proteins from the in-house spectral library acted as the genomic background for statistical testing, and GO terms represented by fewer than five proteins in this library were excluded from the analysis. Significantly enriched GO terms (p < 0.05) were filtered to exclude those with < 1.5-fold enrichment of protein identifications relative to the background library. These were then visualised with Cytoscape 3.6 and manually clustered based on the protein targets shared between GO terms and the semantic similarity measure of Schlicker *et al*.^81^. Node size represents P-value. Node colour represents the median fold-change (vs DMSO) for proteins within each GO term. GO-term clusters that were also represented in an identical analysis of proteins identified in control samples are shaded grey.

A targeted analysis of the 28 proteins within the cell redox homeostasis GO term identified 9 peroxide targets that were enriched by both OZ727 and AA2 within 1 h of treatment (Figure S4A), and alkylation was decreased upon co-incubation of OZ727 with OZ03 or pre-incubation with E64d (Figure S4B). The enriched proteins included several thioredoxin proteins, which are important for maintaining parasite antioxidant defences and regulating redox activity of proteins^53^. Similar trends were seen for these proteins in the OZ747 group (Figure S4A). Oxidation reduction (GO:0055114) and response to oxidative stress (GO:0006979) processes were also significantly enriched biological process GO terms (Supplementary Data 3). Combined, these data suggest that altered redox regulation is an important mechanism for the activity of antimalarial peroxides.

### Validation of recombinant hGrx1-roGFP2 probe with peroxide antimalarial drugs *in vitro*

To further investigate the effects of peroxide antimalarials on the redox status of *P. falciparum* parasites, we used the genetically integrated ratiometric redox-sensor hGrx1-roGFP2 (human glutaredoxin 1 fused to reduction-oxidation sensitive green fluorescent protein), expressed in the cytosol of the NF54*attB P. falciparum* parasite line (NF54*attB*^[hGrx^^1-roGFP2]^)^49^. In order to exclude a direct influence of the drugs on the probe, we initially characterised the interaction of the peroxides with recombinant hGrx1-roGFP2 *in vitro*. DHA, and the ozonides, OZ277 and OZ439, were tested at concentrations ranging from 0.1 µM to 1 mM for up to 10 h in standard reaction buffer containing ferrous iron (to activate the endoperoxide bond) in a plate reader. When compared to DMSO control, the drugs did not significantly affect the fluorescence ratio of recombinant hGrx1-roGFP2 over time (Figure S5), which is consistent with previously published data for peroxides and recombinant hGrx1-roGFP2^54–55^. The recombinant protein was fully oxidised with 1 mM diamide (DIAM) and fully reduced with 10 mM dithiothreitol (DTT) (Figure S5).

### Time-dependent effects of peroxides on the glutathione redox potential of *P. falciparum*

The redox effects of clinically relevant concentrations and exposures of DHA, OZ277 and OZ439 were assessed using NF54*attB*^[hGrx1-roGFP2]^ transgenic parasites in both confocal laser scanning microscopy (CLSM) and plate reader assays (Figure 4, S6A and S6B). To measure the completely oxidised and reduced state of the probe, 30-34 h.p.i trophozoites (6-8% parasitaemia and 2% Hct) were incubated for 5 min with 1 mM DIAM or 10 mM DTT, respectively, blocked with 2 mM NEM and then magnetically enriched. Peroxide-induced effects on parasite oxidative status were monitored at concentrations ranging from 0.1 µM to 1 µM and at intervals between 10 min and 9 h, after which free thiol groups were blocked with 2 mM NEM and trophozoite infected RBCs were magnetically enriched (Figure 4, S6A and S6B).

**Figure 4.**
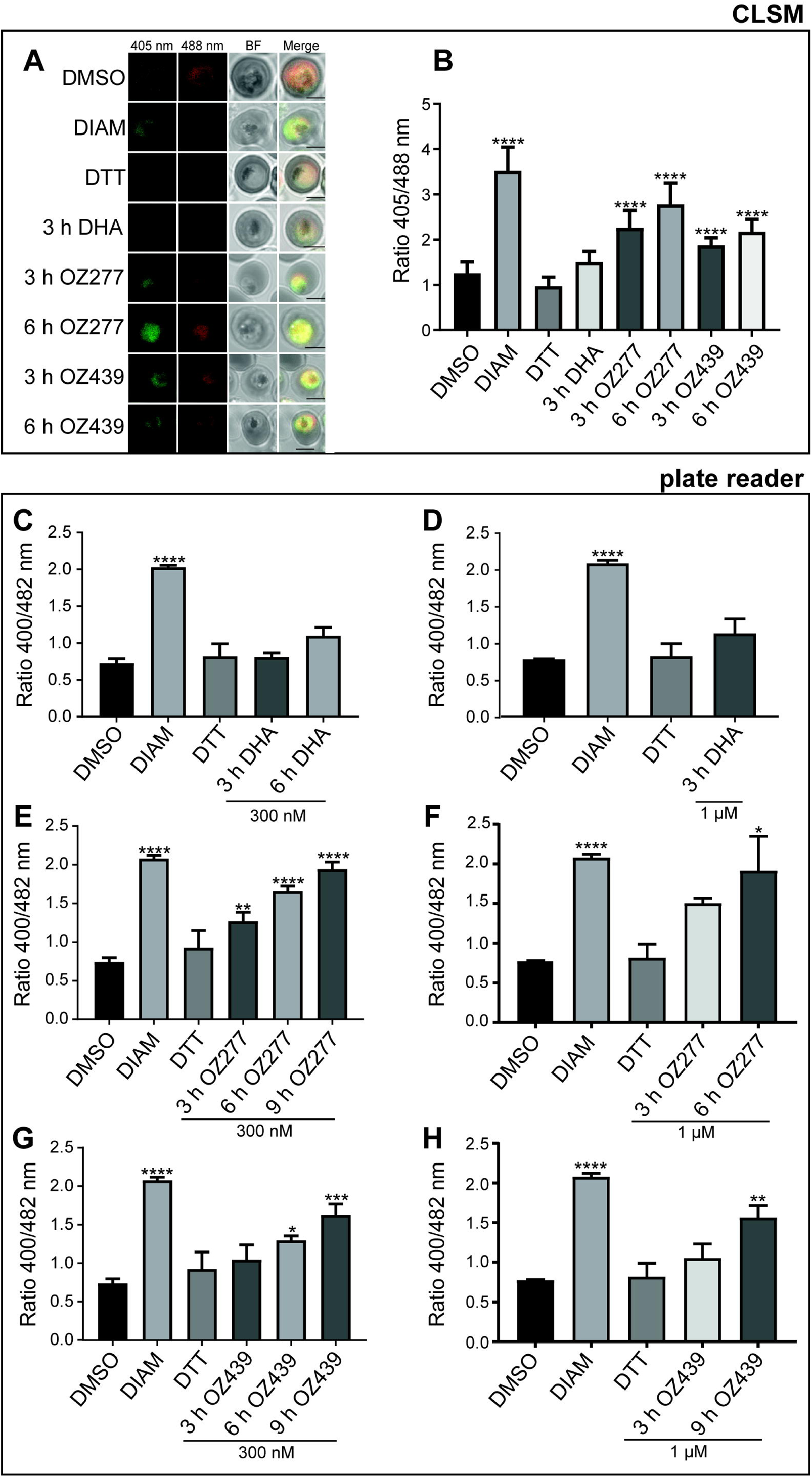
Effects of peroxide antimalarials on the redox ratio of *P. falciparum* NF54*attB*^[hGrx1-roGFP2]^ parasites. NF54*attB*^[hGrx1-roGFP2]^ transgenic parasites were treated with DHA, OZ277 and OZ439 at 300 nM and samples were taken at 3 h post treatment for DHA, OZ277 and OZ439 and at 6 h for OZ277 and OZ439 for the measurement of fluorescence ratio of the redox sensor. Fluorescence ratio of the redox sensor increased for parasites treated with OZ277 and OZ439 for 3 and 6 h treatments, while DHA treatment did not change the redox ratio as measured with confocal laser scanning microscopy (CLSM, A and B). Oxidised (405 nm), reduced (488 nm), BF: bright field, merge: is an overlay of all the panels. The black magnification bar represents 2 µm. Plate reader measurement of the redox ratio following treatment of transgenic parasites with DHA (300 nM (C), 1 µM (D)), OZ277 (300 nM (E), 1 µM (F)) and OZ439 (300 nM (G), 1 µM (H)) showed the same results as CLSM, where only the ozonides affected the redox ratio and this ratio increased over the duration of drug exposure. DMSO-treated parasites at the longest drug incubation duration acted as control, DTT (10 mM, 5 min treatment) was the fully reduced control, while DIAM (1 mM, 5 min treatment) was the fully oxidised control. Columns represent the mean of at least 2 independent experiments with the error bars expressed as SEM. CLSM data were composed of 15-35 trophozoites analysed per experiment for each condition. P-value was calculated using one-way ANOVA, (*, p < 0.05; **, p < 0.01; ***, p < 0.001; ****, p < 0.0001).

Both OZ277 and OZ439 caused time- and concentration-dependent oxidation of the NF54*attB*^[hGrx1-roGFP2]^ parasite cytosol in the CLSM and plate reader when tested at 300 nM for 1-9 h (Figure 4 and S6B). OZ277 at 300 nM increased the fluorescence ratio of NF54*attB*^[hGrx1-roGFP2]^ parasites by 1.27-fold via plate reader (P-value < 0.01) (Figure 4E) and 2.25-fold via CLSM detection (Figure 4A, 4B and S6B) (P-value < 0.0001) within 3 h of drug treatment and oxidation peaked after 9 h (2.0-fold increase, P-value < 0.0001), when compared to the DMSO control. In contrast, OZ439 at 300 nM increased cytosolic oxidation after 6 h of exposure (1.3-fold increase via plate reader detection (P-value < 0.05) (Figure 4G), and 1.8-fold increase via CLSM detection (P-value < 0.0001)) (Figure 4A, 4B and S6B), and had a less pronounced impact compared to OZ277 (2.0-fold) on the fluorescence ratio over a 9 h period (1.63-fold increase, P-value < 0.001), consistent with OZ439 having slower activity kinetics within the parasite^32^. A similar temporal increase in cytosolic oxidation was observed in parasites treated with 1 µM of OZ277 or OZ439 (Figure 4F and H).

Surprisingly, DHA showed no significant effect on the fluorescence ratio of NF54*attB*^[hGrx1-roGFP2]^ parasites in either the CLSM or plate reader when tested at 300 nM for 1, 3 or 6 h (Figure 4A, B, C, D and S6B). To rule out a concentration-dependent effect on the fluorescence ratio and the possibility that DHA induced a rapid oxidative change in the cytosol that quickly returned to basal levels, we also tested DHA concentrations as high as 1 µM (Figure 4D) and exposures as short as 10 min (100 nM) (Figure S6A) in the plate reader assay. No significant change to the fluorescence ratio was detected with short DHA exposures or at a concentration up to 1 µM (Figure 4C and 4D), suggesting that DHA had no effect on the cytosolic glutathione dependent redox ratio of NF54*attB*^[hGrx1-roGFP2]^ parasites under these conditions.

As expected, treatment with 1 mM DIAM led to a significant increase of the fluorescence ratio of NF54*attB*^[hGrx1-roGFP2]^ parasites with both the CLSM and plate reader, whereas there was no change in the fluorescence ratio after treatment with 10 mM DTT, when compared to the untreated control (Figure 4, S6A and S6B). This is consistent with the basal cytosolic environment of *P. falciparum* being strongly reducing^54^.

### Free thiol and glutathione levels in peroxide-treated *P. falciparum* infected RBCs

To further interrogate peroxide-mediated changes to the intracellular redox milieu, we employed a thiol derivatisation targeted mass spectrometry-based metabolomics approach to accurately measure the abundance of small molecule thiols and related metabolites within peroxide-treated infected RBCs (*P. falciparum* 3D7 strain (Figure 5, S7A, and S7B)) (NF54*attB*^[hGrx1-roGFP2]^ strain (Figure S7C)). Trophozoite-infected RBCs (30-34 h.p.i) at 6-8% parasitaemia and 2% Hct were treated with DHA (100 nM for 3 h), OZ277 (300 nM for 3 h) or OZ439 (300 nM for 6 h). Equivalent cultures treated with DIAM (1 mM for 5 min), DTT (10 mM for 5 min) or an equal volume of DMSO (6 h) acted as the oxidation, fully reduced and untreated control, respectively. Following the drug incubation, the cells were magnetically enriched and extracted with 50 mM NEM to rapidly and irreversibly derivatise thiol metabolites and protect them from oxidation during subsequent sample preparation, thus preserving their redox state (method 2) (Figure S7B). As the magnetic enrichment step could alter the cellular redox state prior to NEM stabilisation, we also performed the same drug treatments on magnetically purified cultures that were extracted and derivatised with NEM immediately after the drug incubation (method 3). NEM stabilisation was also performed after drug treatment and prior to the magnet enrichment step (method 1). In all cases, results were compared to DMSO-treated infected RBCs. For peroxide treatments performed on magnetically purified cultures, the activity of the peroxide antimalarials was confirmed to be in the expected nanomolar range for a 3 or 6 h pulse and the IC_50_ values were equivalent in both 3D7 and NF54*attB*^[hGrx1-roGFP2]^ parasites, confirming that the peroxides act similarly in both lines (Figure S7D).

**Figure 5.**
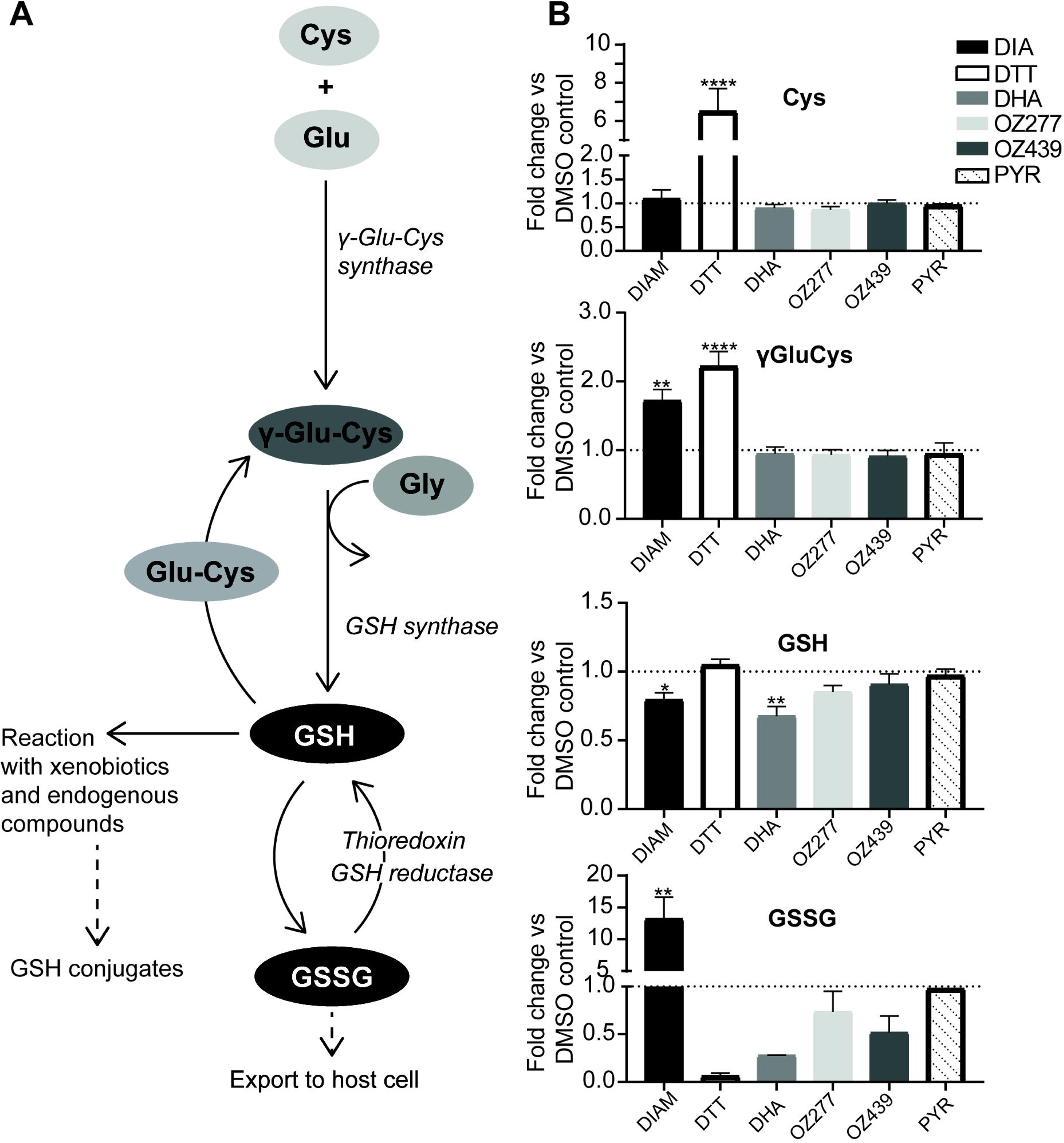
Effect of peroxide antimalarials on thiol levels in *P. falciparum* 3D7 infected RBCs. (A) Schematic of glutathione biosynthesis pathway. (B) Determination of relative levels of NEM-derivatised cysteine (Cys), γ-glutamyl-cysteine (γ-GluCys), reduced glutathione (GSH) and oxidised glutathione (GSSG) compared to DMSO control in *P. falciparum* 3D7 parasites following treatment with DHA (100 nM, 3 h treatment), OZ277 (300 nM, 3 h treatment), OZ439 (300 nM, 6 h treatment) and pyrimethamine (PYR) (1 µM, 5 h treatment) using method 3. DTT (10 mM, 5 min treatment) was the fully reduced control, while DIAM (1 mM, 5 min treatment) was the fully oxidised control. Bars represent the fold change (mean ± SEM) compared to DMSO control. Thiol measurement is from 2-5 biological replicates (with four technical within each biological). P-value was calculated using one-way ANOVA, (*, p < 0.05; **, p < 0.01, p < 0.01; ***, p < 0.0001; ****).

In cultures immediately stabilised with NEM (method 3), the oxidant DIAM significantly depleted GSH (P-value < 0.05) and there was a corresponding increase in the levels of oxidised glutathione (GSSG) (P-value < 0.01), while the reductant DDT depleted GSSG only (Figure 5). This is consistent with DIAM causing an oxidative shift and DTT causing a reducing shift in the free cellular glutathione redox balance and agrees with the fluorescence ratio observed in DIAM- and DTT-treated NF54*attB*^[hGrx1-roGFP2]^ parasites (Figure 4). The glutathione precursor γ-glutamyl cysteine (-GluCys) was increased after DIAM (P-value < 0.01) and DTT (P-value < 0.0001) exposure, likely as a response to restore cellular glutathione levels via increased synthesis (Figure 5). However, cysteine (Cys) levels were not affected by DIAM treatment, but DTT treatment caused a significant increase in Cys levels (P-value < 0.0001) (Figure 5). This significant increase in Cys levels is likely a result of all cystine (the oxidized dimer of Cys) in the infected RBCs being converted to the reduced form (Cys).

Both OZ277 and OZ439 showed a trend towards depleted GSH levels (not significant), with GSSG levels being unaffected (Figure 5). The ozonides caused no detectable changes in Cys, or γ-GluCys levels. DHA treatment depleted the total pool of free glutathione and this included a significant decrease in reduced GSH (P-value < 0.01) and a trend towards decreased oxidised GSSG (1.5- and 3.7-fold decrease, respectively) levels (Figure 5). Cys and γ-GluCys levels were unaffected by DHA treatment (Figure 5). Pyrimethamine, an antimalarial that targets the parasite’s dihydrofolate reductase didn’t affect the relative abundance of thiols in our assay (Figure 5). For most of the compounds tested, similar results were also observed with 3D7 cultures magnetically enriched and stabilised after drug treatment (method 1 and method 2, Figure S7A and B) and in the NF54*attB*^[hGrx1-roGFP2]^ parasite strain (method 3, Figure S7C).

## Discussion

Here, we have used click chemistry based chemical proteomics with an extensive control strategy to show that peroxide antimalarials based on artemisinin and ozonides target similar sets of proteins within *P. falciparum,* and that these proteins are involved in numerous essential biological processes. These findings agree with previous studies demonstrating that ozonides and artemisinins have a similar mode of action and that they disrupt multiple processes important for parasite survival^20–23, 32^.

We also identified a set of proteins that were alkylated exclusively by specific peroxides. This suggests that there may be subtle differences in the underlying mechanisms by which artemisinins, first-generation ozonides and second-generation ozonides kill the malaria parasite, which could have important implications for ozonide cross-resistance with artemisinins.

Functional enrichment analysis of the peroxide antimalarial targets identified food vacuole proteins to be significantly enriched, consistent with this compartment being an important initial site of damage for peroxides^32^, as well as proteins involved in redox processes. Peroxide-induced perturbation of the parasite antioxidant defence system was then confirmed using 1) a parasite cytosolic-specific redox probe, which determined glutathione dependent redox state in real time, and 2) targeted LC-MS based thiol metabolomics, involving derivatisation of thiol metabolites with NEM, to accurately measure total parasite glutathione in both its reduced (GSH) and oxidised (GSSG) forms, and its precursors.

Previous studies using clickable artemisinin and ozonide probes to identify peroxide targets have demonstrated minimal overlap in proteins (25 in total)^21, 23^. When including the targets identified in our study, only 13 proteins were common between all three datasets (Figure S1). Further analysis of these 13 proteins identified them to all be highly abundant in the *P. falciparum* blood-stage proteome (Figure S1), with the exception of PF3D7_0824600 (Fe-S cluster assembly protein DRE2). This minimal overlap could be due to random alkylation of proteins by the peroxides, as proposed by Jourdan *et al*.^20^. However, we identify a significant proportion of proteins from each of these studies individually (36% and 55%), which argues against this hypothesis. An alternative explanation is likely due to the type of mass spectrometry analysis, which in all previous studies utilised the data dependent acquisition approach (DDA). DDA selects precursor ions, corresponding to peptides, for fragmentation in the mass spectrometer based on their abundances, so only the most abundant peptides will produce fragment spectra, which are necessary for protein identification. In a complex sample, this data-dependent approach can lead to inconsistent peptide identification, which limits the number of proteins that can be reproducibly identified and quantified, often favouring highly abundant proteins^56^. In contrast, the mass spectrometry approach used in our study (DIA-MS), selects all precursor ions (peptides) within a mass window and fragments them regardless of their intensity^57^.

This approach results in an extremely high run-to-run reproducibility and a more comprehensive dataset that is not biased towards highly abundant proteins. This is evident from our data, where we have significantly expanded the list of peroxide targets by ∼64 % and the identified proteins ranged in abundance across the *P. falciparum* blood stage proteome (Figure 1). The potential limitation of our DIA approach is the increased identification of low-abundance non-drug-specific proteins that bind to the agarose resin. However, this potential issue was overcome by the inclusion of an extensive set of six control conditions that accounted for any background binding. Overall, the analysis of a total of 36 control samples, alongside the 23 samples from active alkyne-tagged peroxides, provides extremely high confidence in the protein targets identified in this study.

GO analysis of peroxide protein targets identified the food vacuole, and redox homeostasis, as the most significant cellular compartment, and biological process, to be enriched, respectively (Figure 3 and Supplementary Data 2 and 3). Confirming that these proteins are important for the activity of peroxides, alkylation of these proteins with OZ727 was decreased in the presence of the non-clickable parent ozonide, OZ03, and following pre-treatment with the falcipain haemoglobinase inhibitor, E64d, which blocks peroxide activation (Figure S2 and S4). This also confirms that haem derived from haemoglobin digestion is the major iron source responsible for peroxide activation. The food vacuole is a site considered important for haem-dependent activation of peroxides, and protein targets functioning in the food vacuole were also shown to be enriched in previous peroxide mode of action studies^21, 23^. Furthermore, the haemoglobin digestion pathway within the food vacuole is known to be significantly perturbed following treatment with the peroxides^32^.

GO terms associated with general metabolic processes were also specifically enriched in peroxide-treated parasites and not the controls (Figure 3 and Supplementary Data 3). These enriched terms included glycolysis (GO:0006096), which is consistent with previous studies identifying glycolytic proteins as targets of peroxides^21–23^. However, our previous metabolomic analyses revealed the abundance of glycolysis metabolites^32^ and flux through the glycolysis pathway^58^ are unaffected by peroxide exposure. It is possible that a corresponding increase in the abundance of glycolytic proteins^32^ compensates for the loss of enzyme function, raising questions as to the functional importance of these targets to the mechanism of action of peroxides.

The antioxidant system is essential within the parasite, not only for the maintenance of redox homeostasis, but also for the parasite’s ability to respond to drug-induced stress^59^. During blood stage development, reactive oxygen species produced as a by-product of haemoglobin digestion induce intraparasitic oxidative stress, which the parasite manages by engaging a highly efficient antioxidant defence system^59^, which includes glutathione and thioredoxin-related proteins^59^. Parasite glutathione is crucial for maintaining redox balance and has been linked to various drug resistance mechanisms, including resistance against the antimalarial drug chloroquine^59–60^ and more recently artemisinins^61–62^. As glutathione is the major intra-parasitic antioxidant, we used a parasite cytosolic expressed fluorescence-based glutathione biosensor, hGrx1-roGFP2, to monitor glutathione redox potential in infected RBCs treated with various peroxide antimalarials. Interestingly, we found that the ozonides perturbed the parasite cytosolic redox ratio in a concentration- and time-dependent manner, while DHA had no measurable impact on the redox ratio. This was shown using both plate reader and microscopy (Figure 4), and was somewhat surprising considering the involvement of free radicals in the mode of action of artemisinins^63–65^. Interestingly, DHA was shown to increase the redox ratio in a time-dependent manner when the same probe was localised to the parasite mitochondria^55, 61, 66–67^, suggesting that artemisinins differentially affect the redox status of parasite organelles. We therefore, measured total glutathione, and its associated metabolites using a targeted LC-MS based metabolomics method, where we derivatised parasite thiols using NEM to capture the parasite redox state. Our redox analytical method demonstrated that peroxides generally had a trend towards depletion of total glutathione (GSH and GSSG), with DHA having the most profound effect, compared to control (Figure 5). DHA significantly depleted both the reduced (GSH) and oxidised (GSSG) forms of glutathione, which likely explains the lack of impact on the glutathione redox balance (based on the fluorescent biosensor), despite clear depletion of reduced glutathione as measured by LC-MS. In addition to its direct role in redox homeostasis, glutathione plays an important role in repair mechanisms for proteins that have undergone oxidative damage, in addition to potential roles in haem degradation and conjugation of small molecules. Whilst it is possible that glutathione is consumed by direct quenching of the artemisinin-derived radicals^65^, we propose that a significant amount of glutathione is consumed by the parasite’s stress response^32^ to this extensive protein alkylation (Figure 2).

The ozonides demonstrated significantly less depletion of total glutathione levels than DHA, which is consistent with the slower onset of activity of these drugs compared to DHA^13^ and their temporal effects on parasite biochemical pathways^32^. However, the ozonides had a greater impact than DHA on the glutathione redox ratio (Figure 5), and all tested peroxides targeted several proteins involved in redox homeostasis (Figure 3), suggesting that glutathione metabolism plays a critical role in the mode of action of these two classes of peroxide antimalarials, albeit the specific role differs between classes^31, 68–69^. Supporting this hypothesis, pro-oxidant molecules are known to enhance the activity of artemisinins^28^ and parasites with a mutant Kelch 13 propeller allele, linked to artemisinin resistance both *in vitro* and *in vivo*, have elevated glutathione levels^62^. Glutathione has also been associated with artemisinin sensitivity in other parasite lines^70–72^ and we propose that glutathione metabolism may provide opportunities for intervention with pro-oxidant combinations to enhance peroxide sensitivity in the context of artemisinin resistance. ^73^

We have demonstrated with chemical proteomics that peroxide antimalarials alkylate a range of *Plasmodium* proteins that are vital for parasite growth and survival. The parasite antioxidant system was one of the crucial pathways targeted. Using genetically-encoded redox sensors and LCMS-based targeted thiol measurements, we showed that peroxide antimalarials cause significant disruption to *P. falciparum* glutathione metabolism. This detailed description of the protein targets of artemisinins and ozonides, and the important role of redox metabolism in their modes of action, will underpin future research to optimise their efficacy against increasingly common artemisinin resistant parasites.

## Methods

### P. falciparum cell culture

3D7 and the NF54*attB*^[hGrx1-roGFP2]^ *P. falciparum* transgenic line^49^ were cultured as previously described^74^. Parasites were tightly synchronised by double sorbitol lysis 14 h apart.

### Peroxide antimalarials and alkyne tagged peroxide probes

DHA and artemisinin were purchased from Jomar Bioscience and Sigma-Aldrich, respectively. OZ277 and OZ439 were provided by the Medicines for Malaria Venture (Geneva). OZ03, the artemisinin alkyne, AA2, and ozonide alkynes, OZ727 and carbaOZ727, were obtained by previously published procedures^20–21, 75^. Synthesis of the alkyne ozonide OZ747, the click chemistry probe analogous to the advanced-stage antimalarial clinical candidate, OZ439 (artefenomel), is described below.

### Synthesis of OZ747

#### *cis-*6-(Pent-4-ynamido)adamantane-2-spiro-3’-8’-[4-(2-morpholinoethoxy)phenyl]-1’,2’,4’-trioxaspiro[4.5]decane mesylate (OZ747)

**Step 1.** To a solution of *cis-*6-oxoadamantane-2-spiro-3’-8’-(4-hydroxyphenyl)-1’,2’,4’-trioxaspiro[4.5]decane^76^ (500 mg, 1.35 mmol) in dry DME (25 mL) were added powder NaOH (324 mg, 8.01 mmol) and Bu_4_NHSO_4_ (93 mg, 0.50 mmol). The resulting mixture was stirred at rt for 30 min before addition of *N*-(2-chloroethyl)morpholine hydrochloride (415 mg, 2.22 mmol). The resulting reaction mixture was stirred at 60 °C for 12 h. After filtration of the solid material, the filtrate was concentrated in vacuo to afford a residue which was dissolved in EA (100 mL), washed with H_2_O (50 mL) and brine (50 mL), dried over MgSO_4_, and filtered and concentrated in vacuo to afford *cis-*6-oxoadamantane-2-spiro-3’-8’-[4-(2-morpholinoethoxy)phenyl]-1’,2’,4’-trioxaspiro[4.5]decane (560 mg, 86%) as a white solid. mp 145 – 146 °C. ^1^H NMR (500 MHz, CDCl_3_) δ 1.67 – 1.75 (m, 2H), 1.84 – 1.92 (m, 4H), 1.94 (d, *J* = 13.0 Hz, 2H), 1.99 (d, *J* = 13.0 Hz, 2H), 2.06 (d, *J* = 13.0 Hz, 2H), 2.15 (s, 2H), 2.26 (d, *J* = 13.0 Hz, 2H), 2.35 (d, *J* = 13.0 Hz, 2H), 2.50 (m, 3H), 2.57 (s, 4H), 2.79 (t, *J* = 5.5 Hz, 2H), 3.73 (t, *J* = 4.5 Hz, 4H), 4.09 (t, *J* = 5.5 Hz, 2H), 6.84 (d, *J* = 8.5 Hz, 2H), 7.12 (d, *J* = 8.5 Hz, 2H); ^13^C NMR (125 MHz, CDCl_3_) δ 31.57, 34.62, 35.68, 35.75, 35.90, 41.97, 44.72, 45.15, 54.09, 57.70, 65.82, 66.95, 109.04, 108.28, 114.53, 127.62, 138.30, 157.16, 215.88. Anal. calcd for C_28_H_37_NO_6_: C, 69.54; H, 7.71; N, 2.90. Found: C, 70.00; H, 7.47; N, 2.77. **Step 2.** Following the method of Wu et al.^76^, a one-pot reductive amination/acylation of *cis-*6-oxoadamantane-2-spiro-3’-8’-[4-(2-morpholinoethoxy)phenyl]-1’,2’,4’-trioxaspiro[4.5]decane (430 mg, 0.89 mmol) afforded the free base of **OZ747** (170 mg, 34%). **Step 3.** To a solution of the free base of **OZ747** (170 mg, 0.30 mmol) in EA (5 mL) was added dropwise a solution of methanesulfonic acid (58 mg, 0.60 mmol) in diethyl ether (1 mL). The mixture was stirred at rt for 30 min, and the resulting solid was filtered, washed with cold EA, and dried in vacuo at 50 °C to afford **OZ747** as a white solid (157 mg, 79%). mp 151 – 152 °C. ^1^H NMR (500 MHz, DMSO-*d*_6_) δ 9.91 (s, 1H), 7.84 (d, *J* = 7.3 Hz, 1H), 7.17 (d, *J* = 8.4 Hz, 2H), 6.93 (d, *J* = 8.3 Hz, 2H), 4.32 (t, *J* = 4.7 Hz, 2H), 3.98 (dd, *J* = 12.9, 3.4 Hz, 2H), 3.77 (d, *J* = 7.0 Hz, 1H), 3.71 (t, *J* = 12.5 Hz, 2H), 3.61 – 3.53 (m, 2H), 3.50 (d, *J* = 12.5 Hz, 2H), 3.20 (q, *J* = 11.5 Hz, 2H), 2.75 (s, 1H), 2.57 (t, *J* = 11.9 Hz, 1H), 2.40 – 2.24 (m, 7H), 2.09 – 1.62 (m, 17H), 1.55 (m, 3H); ^13^C NMR (125 MHz, DMSO-*d*_6_) 170.39, 156.30, 139.39, 128.04, δ 115.16, 110.74, 108.70, 84.24, 71.74, 63.70, 62.43, 55.57, 52.48, 52.18, 41.17, 40.24, 35.50, 35.35, 34.54, 34.51, 34.20, 31.75, 30.62, 30.22, 28.83, 28.78, 14.93. Anal. calcd for C_34_H_48_N_2_O_9_S: C, 61.80; H, 7.32; N, 4.24. Found: C, 61.56; H, 7.10; N 4.00.

### Drug sensitivity assays

Parasite growth inhibition assays with the peroxide antimalarials and the clickable alkyne tagged peroxide probes were performed as previously described^12^. Briefly, for clickable alkyne tagged peroxides (OZ727, OZ747, AA2 and carbaOZ727) and their associated non-clickable controls (OZ03, OZ277, OZ439, DHA and artemisinin), IC_50_s were performed on *P. falciparum* (3D7) infected cultures at 1% parasitaemia and 2% haematocrit (Hct) in a 96 well plate over 48 h at 37 °C. For the non-clickable peroxides, OZ277, OZ439 and DHA, drug pulse assays were performed on both 3D7 and NF54*attB*^[hGrx1-roGFP2]^ *P. falciparum* lines that were magnet harvested (90% parasitemia) and adjusted to a final Hct of 0.12%. Following a drug pulse of 3 h for DHA and OZ277 and 6 h for OZ439, peroxides were washed off as previously described^12^ with minor modification. Following four washes with 200 µL of complete RPMI culture media containing 5% Albumax II, an extra wash with complete RPMI media was also included. After the washes, parasitaemia was adjusted to 0.6% and the Hct was returned to 2% using uninfected RBCs. Cultures were then transferred to a flat-bottom 96 well microplate and incubated at 37 °C for 48 h. Following the 48 h incubation, parasite drug susceptibility was assessed by measurement of SYBR green I fluorescence as previously described^12^.

The data was analysed using GraphPad Prism software version 8.0.2 as previously described^12^.

### Parasite treatment for chemical proteomics

*P. falciparum* parasites (3D7) synchronised to the trophozoite stage (28-32 h post invasion (h.p.i)) were diluted to 10% parasitaemia and adjusted to 2% Hct. For each treatment condition, 30 mL of infected RBC culture was used. Cultures were treated with 300 nM of the active clickable peroxides AA2, OZ727 or OZ747 for 1 to 6 h. Control cultures were incubated with 300 nM of the inactive (clickable) non-peroxide probe, carbaOZ727 (1 h), 300 nM of the non-clickable parent ozonide, OZ03 (1, 3 and 6 h), or an equivalent volume of DMSO (1 h, < 0.01% final concentration). For competition experiments, the cultures were treated together with 300 nM of OZ727 and either an equivalent (300 nM) or excessive (900 nM) concentration of OZ03 for 1 h. In experiments using the cysteine protease inhibitor E64d to antagonise peroxide activity, cultures were pre-treated for 30 min with 10 µM of E64d, followed by incubation with OZ727 for an additional 1 h. For all experiments, cultures were maintained at 37 °C under a gas atmosphere of 94% N_2_, 5% CO_2_ and 1% O_2_ for the duration of drug exposure.

After incubation with the alkyne probes, all subsequent steps were performed on ice or at 4 °C. The cultures were centrifuged at 700 g for 3 min to remove the media and parasites were isolated from the RBC by incubating with 0.1% saponin in PBS containing protease (Roche) and phosphatase inhibitors (Sigma, 20 mM sodium fluoride, 0.1mM sodium orthovanadate and 10 mM beta glycerophosphate) for 10 min. The cell lysates were then centrifuged at 2,576 x *g* to remove RBC proteins and the resulting parasite pellets were washed a further three times in PBS containing protease and phosphatase inhibitors to ensure removal of RBC membrane debris. Parasite pellets were stored at −80 °C until protein extraction and click chemistry enrichment of probe labelled proteins.

### Copper catalysed click chemistry enrichment of alkylated proteins

Proteins were extracted by solubilising parasite proteins using 0.5% sodium deoxycholate (SDC) in 100 mM HEPES buffer and heating each sample at 90 °C for 5 min. This was followed by 3 x 30 sec cycles of probe sonication on ice. The soluble fraction was separated by centrifugation at 14,800 x *g* for 5 min and the protein concentration determined using the Pierce BCA assay kit (Thermo Scientific™ Pierce™). The protein concentration of each sample was adjusted to 693 µg/mL. Alkyne labelled proteins were enriched and affinity purified by direct attachment to azide agarose beads (Jena Bioscience) using copper catalysed click chemistry. For each reaction, 50 µL of azide agarose beads (prewashed with 1,400 µL of Milli-Q water), 22.6 µL of TCEP (50 mM in Milli-Q water), 68 µL of THPTA ligand (1.7 mM in DMSO/*t*-butanol, 1:4 ratio) and 22.6 µL of CuSO4 (50 mM in Milli-Q water) were sequentially added to 837 µL of the cell lysate. The lysate was incubated at room temperature for 1 h while rotating end-over-end. Following the click reaction, the agarose resins containing the clicked proteins were washed with 1 mL of Milli-Q water.

### Washing and on-bead digestion of clicked proteins

The washed resins with the clicked proteins attached were resuspended in 1 mL of agarose wash buffer (100 mM HEPES, 1% SDS, 250 mM NaCl, 5 mM EDTA, pH 8). Samples were reduced by addition of 10 µL of 1 M DTT and heated at 70 °C for 15 min, before being allowed to cool for a further 15 min. The supernatant was then removed by centrifugation at 4000 g for 5 min and the samples were alkylated in the dark with 40 mM iodoacetamide in 1 mL of agarose wash buffer for 30 min at room temperature. The beads were then extensively washed five times with agarose wash buffer, five times with 8 M urea and a further 5 times with 20% acetonitrile (ACN). Between each wash, samples were centrifuged at 4000 g for 3 min. Following the final wash, the resins were resuspended in 500 µL of digestion buffer (100 mM HEPES, 2 mM CaCl_2_, 10% ACN) and the contents were transferred to a fresh tube. The tube was rinsed with a further 500 µL of digestion buffer and this was transferred to the same fresh tube. The resins were then pelleted by centrifugation at 4000 g for 5 min and ∼800 µL of supernatant was removed. On-bead protein digestion was performed by adding 1 µg of sequencing grade trypsin (Promega) to the samples, which were incubated overnight at 37 °C while spinning end-over-end. On the following day, the beads were separated from the digested peptides by centrifuging the samples at 4000 g for 5 min and supernatant transferred to a clean tube. The resins were briefly vortexed and washed with a further 500 µL of Milli-Q water, which was then transferred to the same clean tube containing the digested peptides. Each sample was made up to 1 mL volume with Milli-Q water and acidified by addition of 2 µL of formic acid. The samples were then desalted using in-house generated StageTips^77^, dried and resuspended in 10 µL of 2% ACN and 0.1% formic acid containing indexed retention time (RT) peptides for LCMS/MS analysis.

### Liquid chromatography mass spectrometry (LC-MS/MS) analysis

Proteomics LC-MS/MS data was acquired using Dionex Ultimate® 3000 RSLCnano system coupled to Q Exactive HF mass spectrometer (Thermo Scientific) carried out as described previously ^78^, with minor modifications. MS2 data was collected in data independent mode (DIA) with a 33-fixed window setup of 18 *m/z* effective precursor isolation over the *m/z* range of 375-975 Da.

### Chemical proteomics data analysis

Raw files were processed using Spectronaut^TM^ 13.0 against an in-house generated asexual *P. falciparum* spectral library. The library contained 44 449 peptides corresponding to 4730 proteins, of which 3113 were *P. falciparum* proteins^78^. For processing, raw files were loaded and Spectronaut calculated the ideal mass tolerances for data extraction and scoring based on extensive mass calibration using a correction factor of one. Both at the precursor and fragment levels, the highest datapoint within the selected *m/z* tolerance was chosen for targeted data extraction. Identification of peptides against the library was based on the default Spectronaut settings (Manual for Spectronaut 13.0, available on Biognosis website). Briefly, the Q-value cut-off at the precursor and protein levels were set at 1%, therefore only those that passed this threshold were considered as identified and used for other subsequent processes. RT prediction type was set to dynamic indexed RT. Interference correction was on MS2 level. For quantification, the interference correction was activated and a significance filter of 0.01 was used for Q-value filtering with the Q-value sparse setting applied. The imputing strategy was set to no imputing.

Identified proteins in the OZ727 and OZ747 groups with a fold-change ≥ 2 compared to the DMSO, carbaOZ727 and time-matched OZ03 controls, and with a p-value < 0.05 (Mann-Whitney *U* test) compared to carbaOZ727, were considered as protein targets. The same thresholds were applied to proteins in the AA2 group, except p-value filtering was based on comparison with the DMSO control.

The identified protein targets for OZ727, OZ747 and AA2 were subjected to gene ontology (GO) enrichment analysis with topGO^79^. PlasmoDB GO terms were used for GO terms mapping and the elim algorithm was applied to navigate the topology of the GO graph^80^. The Fisher exact test was used for testing enrichment of GO terms. *P. falciparum* proteins from the in-house spectral library were used as the genomic background for statistical testing, and GO terms represented by fewer than five proteins in this library were excluded from the analysis. Significantly enriched GO terms (p < 0.05 and enrichment > 1.5-fold) were visualised in Cytoscape 3.6 and manually clustered based on the protein targets shared between GO terms and the semantic similarity measure of Schlicker *et al*.^81^.

The mass spectrometry proteomics data have been deposited to the ProteomeXchange Consortium via the PRIDE^82^ partner repository with the data set identifier PXD027334. [Reviewer account details are username: and password: 5XhN5Wjo]

### *In vitro* interaction of peroxide antimalarials with the hGrx1-roGFP2 recombinant protein

*In vitro* measurements with the recombinant hGrx1-roGFP2 redox sensor were carried out as previously described^49, 54–55^. Peroxides, diamide (DIAM) and dithiothreitol (DTT) were diluted with a degassed reaction buffer (100 mM potassium phosphate, 1 mM ethylenedaiminetetraacetic acid, EDTA, pH 7.0) and used immediately. Purified recombinant protein was prepared as previously described^49, 55^ and diluted in reaction buffer to a concentration of 1.25 µM, so that in the plate assay, the final concentration is 1 µM. Peroxides at different concentrations combined with DIAM and DTT in the presence of iron (II) chloride (Sigma) (8.7 mM final concentration) were mixed with hGrx1-roGFP2 in a 96-well plate (black, µClear TC Greiner Bio-One). The emission at 510 nm after excitation at 405 and 480 nm was measured after 5 min, 30 min, 1, 3, 5, and 10 h in a plate reader with optimal reading settings. Data from two independent experiments with two technical replicates were analysed for each concentration.

### Effects of the peroxides and other antimalarial drugs on redox homeostasis

The effect of peroxides on *P. falciparum* redox homeostasis was investigated on late trophozoites (30-34 h.p.i) using the NF54*attB*^[hGrx1-roGFP2]^ parasite line and were carried out as previously described^49^. Briefly, parasites (8% parasiteamia, 2% Hct) were incubated with 0.1 to 1 µM of each peroxide antimalarial for 10 min to 9 h, depending on the compound. Following the appropriate incubation period, free thiol groups were blocked with 2 mM *N*-ethylmaleimide (NEM) for 15 min at 37 °C.

After incubation with NEM, the infected RBC cultures were washed and resuspended in Ringer’s solution. Resuspended cells were then magnetically enriched (80% parasiteamia) (Miltenyi Biotech, Bergisch Gladbach, Germany) and the parasite redox ratio was measured using either the confocal live-cell imaging or plate reader analyses. All peroxide incubation experiments included an untreated control group (“mock-treated” with DMSO for 9 h), as well as controls for complete oxidation (1 mM DIAM for 5 min) and complete reduction (10 mM DTT for 5 min).

### Confocal live-cell imaging and image processing

Live-cell imaging was performed as previously described^49^. Briefly, drug-treated cells that had been magnet harvested and resuspended in pre-warmed Ringer’s solution were seeded onto poly-_L_-lysine-coated flat µ-slides VI (Ibidi, Martinsried, Germany). The probes were excited at 405 and 488 nm and emissions were detected at 500-550 nm. Images were analysed with a custom written macro in the Fiji distribution of ImageJ^83^. The macro was as follows: Region of interest (ROIs) of the cells were generated by applying a gaussian blur (sigma=4), generating a binary image and applying a binary watershed. 32-bit ratio images were generated by dividing the two fluorescent channels. The ROIs were used to measure the fluorescent intensity ratio in each cell. Images of the analysed cells were saved out and manually curated to select only infected RBCs that showed fluorescent signals at both 405 and 488 nm excitation and had an intact host cell. The 405/488 nm ratios were calculated and graphs were plotted using GraphPad Prism 8.0.2 software. One-way ANOVA was applied for statistical analysis of significance (****, p < 0.0001).

### Plate reader analyses of redox homeostasis

Plate reader analyses were performed as previously described^49^. Briefly, magnet harvested drug-treated NF54*attB*^[hGrx1-roGFP2]^ parasites (80% parasiteamia) with their free thiols blocked by NEM were counted using the improved Neubauer hemocytometer (Brand GmbH, Wertheim, Germany) and adjusted to 2.0 x 10^5^ iRBCs/µL. Parasites (20 µL) were transferred to 384-well plate and emission (530 nm) was measured with excitation wavelengths of 400 and 482 nm using a Clariostar plate reader (multi-chromatic) (BMG Labtech, Ortenberg, Germany). The calculated ratio of 400/482 nm was then plotted using GraphPad Prism 8.0.2 software. One-way ANOVA was applied for statistical analysis of significance (*, p < 0.05; **, p < 0.01; ***, p < 0.001; ****, p < 0.0001).

### LC-MS determination of free thiol and glutathione levels in peroxide-treated P. falciparum infected RBCs

Levels of reduced glutathione (GSH) and other thiol metabolites were determined by LC-MS for 3D7 and NF54*attB*^[hGrx1-roGFP2]^ *P. falciparum* parasites following treatment with peroxides. Samples were prepared via several different methodologies to obtain the most reproducible output and ensure that no major experimental artefacts were introduced during the sample processing. For simplicity, these are referred as method 1, 2 and 3. The results from method 3 are reported in the main manuscript (Figure 5), while the results from methods 1 and 2 are shown in the supplementary material (Figure S7).

Method 1 sample preparation was the same as experiments used to measure the parasite redox ratio. Briefly, infected RBC cultures were incubated with the test compounds for the desired period (100 nM DHA for 3 h, 300 nM OZ277 for 3 h, 300 nM OZ439 for 6 h, 1 mM DIAM for 5 min and DMSO for 6 h) and thiols were quenched by adding freshly prepared NEM solution to the final concentration of 2 mM in culture media. The cultures were then magnet harvested and the parasitemia for each condition was adjusted to be 80%. The cells (5 x 10^7^) were pelleted by centrifugation for 5 min at 1000 g and the supernatant was removed. The cells were then washed with PBS, prior to addition of 100 µL ice cold methanol for extraction of thiols.

For method 2 sample preparation, the initial sample preparation was the same as described for method 1, except that NEM was added to the extraction solvent (50 µL of 50 mM NEM in 80% methanol, 20% 10 mM ammonium formate, pH 7.0 and 50 µL of 100% ACN). Therefore, free thiols were derivatised at the extraction step.

For method 3, magnetically enriched cultures at 90% parasitaemia and 0.12% Hct were incubated with the test compounds for the desired times (drug treatments, 100 nM DHA for 3 h, 300 nM OZ277 for 3 h, 300 nM OZ439 for 6 h, 1 µM pyrimethamine for 5 h, and three control conditions, 1 mM DIAM for 5 min, 10 mM DTT for 5 min and DMSO for 6 h). Following the drug incubation, cells were washed with PBS and 5 x 10^7^ cells were used for thiol metabolite extraction. Thiols were extracted using NEM-containing extraction solvent (50 µL of 50 mM NEM in 80% methanol, 20% 10 mM ammonium formate, pH 7.0 and 50 µL of 100% ACN).

For all methods, following thiol extraction with the appropriate solvent, cells were incubated on a shaker for 1 h at 4 °C and then centrifuged at 20,000 g to remove insoluble material. Supernatants were transferred to glass HPLC vials and stored at −80 °C until analysis.

### LC-MS thiol analysis and data processing

LC-MS data was acquired on a Q-Exactive Orbitrap mass spectrometer (Thermo Scientific, Waltham, Massachusetts, United States) coupled with a Dionex Ultimate® 3000 RS (Thermo Scientific) HPLC system. The method was adapted from previously described procedure^84, 85^ with some changes. Chromatographic separation was performed on Poroshel Infinity HILIC-Z column (2.7 µm, 2.1 × 100 mm, Agilent Technologies) kept at 25°C. The total run time was 20 min at 0.3 ml/min flowrate with an injection volume of 10 µL. Buffer A was 20 mM ammonium carbonate, buffer B was acetonitrile, syringe wash solvent was 50% isopropanol. Gradient started 90% B, decreased to 65% B at 10 min, further decreased to 20% B at 11.5 min, kept at 20% B until 13 min, returned to 90% B at 14 min and kept at 90% B until 20 min. HESI ion source parameters were as follows: sheath gas 50, aux gas 20 and sweep gas 2 arbitrary units, spray voltage was set to 4.0 kV, capillary temperature to 300 °C, S-Lens RF level to 50 and auxiliary gas heater temperature to 120 °C. The mass spectrometer was operated in full scan mode in positive detection mode at 35 000 resolution at 200 m/z with detection range of 85 to 1275 m/z. The samples were analysed as a single batch to reduce batch-to-batch variation and randomised to account for LC-MS system drift over time.

Data was processed using peak areas, which were obtained with QuanBrowser (Thermo Scientific) software, version 4.2 by integrating MS1 peaks. Peak identities were then confirmed using NEM derivatised authentic standards. The calculated peak area was plotted using GraphPad Prism 8.0.2 software and one-way ANOVA was applied for statistical analysis (*, p < 0.05; **, p < 0.01; ***, p < 0.001; ****, p < 0.0001).

## Ancillary Information

### Supporting Information

Comparison of pulldown results with previously published datasets; heatmap representation of the effect of OZ03 co-incubation or E64d pre-treatment on OZ727 labelling of food vacuole proteins (GO:0020020); GO biological process enrichment analysis of control dataset; alkylation of redox homeostasis proteins (GO:0045454) by peroxide probes and effect of OZ03 co-incubation or E64d pre-treatment on OZ727 labelling of redox homeostasis proteins; effects of peroxide antimalarial on reduced hGrx1-roGFP2; effect of peroxide antimalarials on the redox ratio of *P. falciparum* NF54*attB*^[hGrx1-roGFP2]^ parasites; effect on thiols (using different sample preparation methods) and activity of peroxide antimalarials in *P. falciparum* 3D7 and NF54*attB*^[hGrx1-roGFP2]^ parasites Supplementary Data 1. Complete proteomics dataset for identification of protein targets of peroxide antimalarials and heatmap representation of all protein binding targets of OZ727, OZ747 and AA2 following treatment of *P. falciparum* infected RBCs for 1 hr (all 300 nM) with protein IDs and significant proteins (+) Supplementary Data 2. GO enrichment analysis results for cellular component Supplementary Data 3. GO enrichment analysis results for biological process

## Acknowledgment

Funding support was provided by the NHMRC (APP#1160705, APP1128003, APP1185354 and APP1148700) and the NIH (AI116723-01). The authors acknowledge the Monash Proteomics and Metabolomics Facility (Parkville Node) for providing LC-MS technical assistance. The Australian Red Cross Blood Service in Melbourne donated human red blood cells for *in vitro* parasite cultivation.

## Author Contributions

DJC, KB, GS and CG directed the overall research program; GS, CG, ADP, AKS, KCH, DA performed the experiments; GS, CG, ADP, TGB, and CAM analysed the data and prepared figures; JW, XW, YD and JLV made the click chemistry probes, all authors contributed to experimental design and wrote and edited the manuscript.

## Abbreviations Used

ACN, acetonitrile; ACTs, artemisinin-based combination therapies; Cys, cysteine; DHA, dihydroartemisinin; DIAM, diamide; DIA-MS, data independent acquisition mass spectrometry; DDA, data dependent analysis, DMSO, dimethylsulfoxide; DTT, Dithiothreitol; GO, gene ontology; GSH, reduced glutathione; GSSG, oxidised glutathione; h, hour; Hct, haematocrit; hGrx-1-roGFP2, glutaredoxin 1 fused to a reduction-oxidation sensitive GFP; IC_50,_ 50% inhibition concentration; iRT, indexed retention time; LC-MS, liquid chromatography-mass spectrometry, m/z, mass to charge ratio; mM, millimolar; NEM, *N*-ethylmaleimide; PBS, phosphate-buffered saline; *P. falciparum, Plasmodium falciparum*; RBCs, red blood cells; SDC, sodium deoxycholate; γ-GluCys, γ-glutamyl cysteine; µL, microlitre; µM, micromolar.

