## Supplemental Figures for "Peroxide antimalarial drugs target redox homeostasis in *Plasmodium falciparum* infected red blood cells"

A

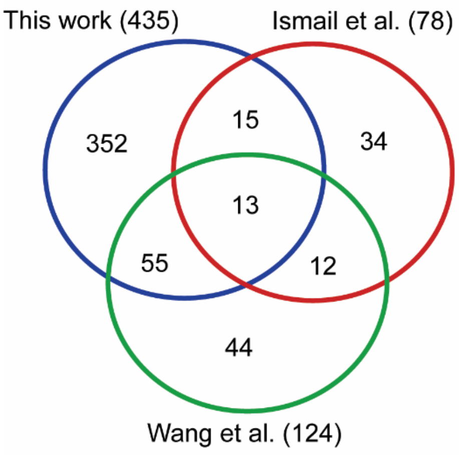


| Enriched proteins that are common between this work (Siddiqui et al.), Wang et al., and Ismail., et al. | |
|---|---|
| **Gene ID** | **Protein description** |
| PF3D7_1015900 | Enolase (ENO) |
| PF3D7_1246200 | Actin I (ACT1) |
| PF3D7_0608800 | Ornithine aminotransferase (OAT) |
| PF3D7_0523000 | Multidrug resistance protein (MDR1) |
| PF3D7_1324900 | L-lactate dehydrogenase (LDH) |
| PF3D7_0322900 | 40S ribosomal protein S3A, putative (S3A) |
| PF3D7_1408000 | Plasmepsin II (PM II) |
| PF3D7_0903700 | α tubulin 1 |
| PF3D7_0709000 | Chloroquine resistance transporter (CRT) |
| PF3D7_0102200 | Ring-infected erythrocyte surface antigen (RESA) |
| PF3D7_0207600 | Serine repeat antigen 5 (SERA5) |
| PF3D7_0930300 | Merozoite surface protein 1 (MSP1) |
| PF3D7_0824600 | Fe-S cluster assembly protein DRE2, putative (DRE2) |

B

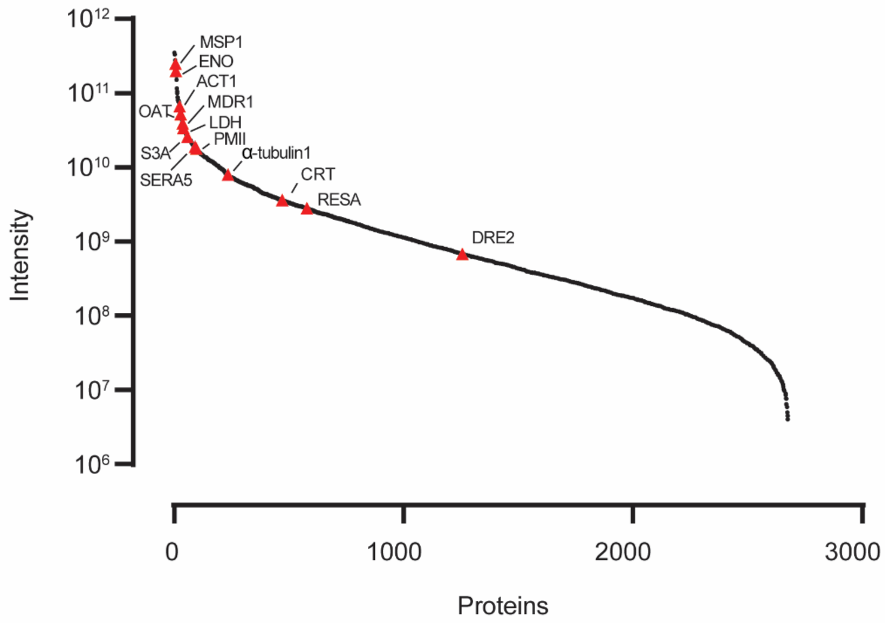


**Figure S1. Comparison of pulldown results with previously published datasets.** (A) Venn diagram showing the overlap between proteins enriched with peroxide clickable probes in the current study (Siddiqui *et al.*), Ismail *et al.*[1] and Wang *et al.*[2] and table with the 13 commonly identified proteins. (B) Scatterplot showing that the 13 commonly identified proteins (red triangles) from each of the three studies are generally highly expressed proteins within the *P. falciparum* blood stage proteome, except DRE2. The relative abundance of proteins in the *P. falciparum* blood stage proteome (Siddiqui *et al.*, unpublished data) are shown as black circles.

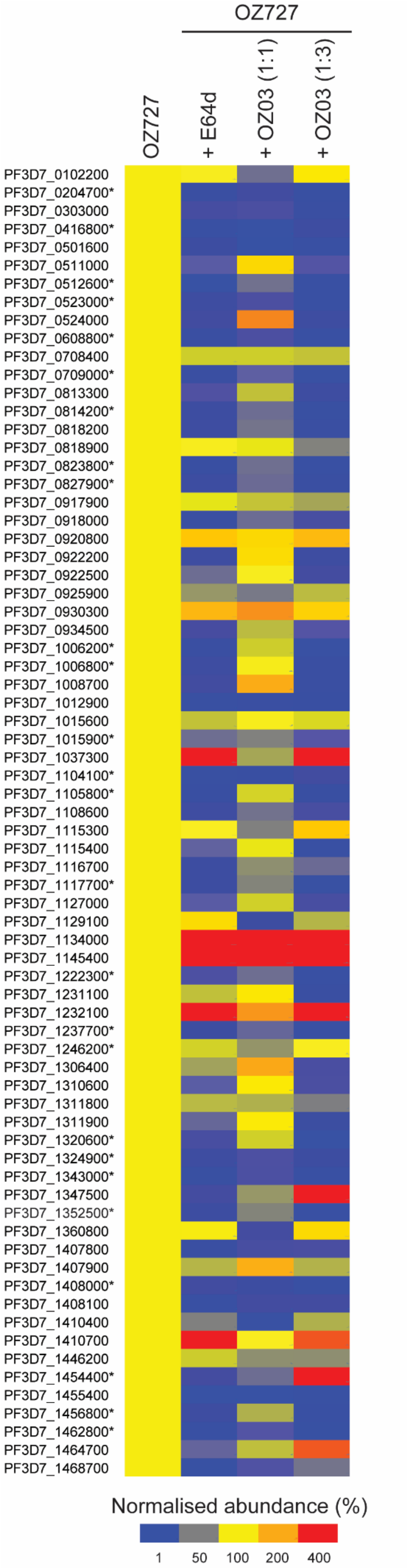

OZ727
OZ727
+ E64d
+ OZ03 (1:1)
+ OZ03 (1:3)
PF3D7_0102200
PF3D7_0204700*
PF3D7_0303000
PF3D7_0416800*
PF3D7_0501600
PF3D7_0511000
PF3D7_0512600*
PF3D7_0523000*
PF3D7_0524000
PF3D7_0608800*
PF3D7_0708400
PF3D7_0709000*
PF3D7_0813300
PF3D7_0814200*
PF3D7_0818200
PF3D7_0818900
PF3D7_0823800*
PF3D7_0827900*
PF3D7_0917900
PF3D7_0918000
PF3D7_0920800
PF3D7_0922200
PF3D7_0922500
PF3D7_0925900
PF3D7_0930300
PF3D7_0934500
PF3D7_1006200*
PF3D7_1006800*
PF3D7_1008700
PF3D7_1012900
PF3D7_1015600
PF3D7_1015900*
PF3D7_1037300
PF3D7_1104100*
PF3D7_1105800*
PF3D7_1108600
PF3D7_1115300
PF3D7_1115400
PF3D7_1116700
PF3D7_1117700*
PF3D7_1127000
PF3D7_1129100
PF3D7_1134000
PF3D7_1145400
PF3D7_1222300*
PF3D7_1231100
PF3D7_1232100
PF3D7_1237700*
PF3D7_1246200*
PF3D7_1306400
PF3D7_1310600
PF3D7_1311800
PF3D7_1311900
PF3D7_1320600*
PF3D7_1324900*
PF3D7_1343000*
PF3D7_1347500
PF3D7_1352500*
PF3D7_1360800
PF3D7_1407800
PF3D7_1407900
PF3D7_1408000*
PF3D7_1408100
PF3D7_1410400
PF3D7_1410700
PF3D7_1446200
PF3D7_1454400*
PF3D7_1455400
PF3D7_1456800*
PF3D7_1462800*
PF3D7_1464700
PF3D7_1468700
Normalised abundance (%)
1
50
100
200
400

**Figure S2. Heatmap representation of the effect of OZ03 co-incubation or E64d pre-treatment on OZ727 labelling of food vacuole proteins (GO:0020020).** The extent of food vacuole protein (GO:0020020) alkylation is expressed relative to levels obtained after treatment with 300 nM of OZ727 alone for 1 h (column 1). OZ727 activation was blocked by pre-treatment (30 min) of live parasite cultures with 10 µM of the cysteine protease inhibitor, E64d, followed by incubation with OZ727 for an additional 1 h (column 2). Competition of OZ727 binding to target proteins, was achieved by co-incubation with 300 nM of OZ727 and either an equivalent (300 nM, column 3) or excessive (900 nM, column 4) concentration of OZ03 for 1 h. The average protein abundance from at least three biological replicates is presented. Yellow features represent no change compared to OZ727 alone. Increased and decreased levels are represented by red and blue, respectively. Significantly enriched food vacuole proteins identified as OZ727 targets are indicated (*).

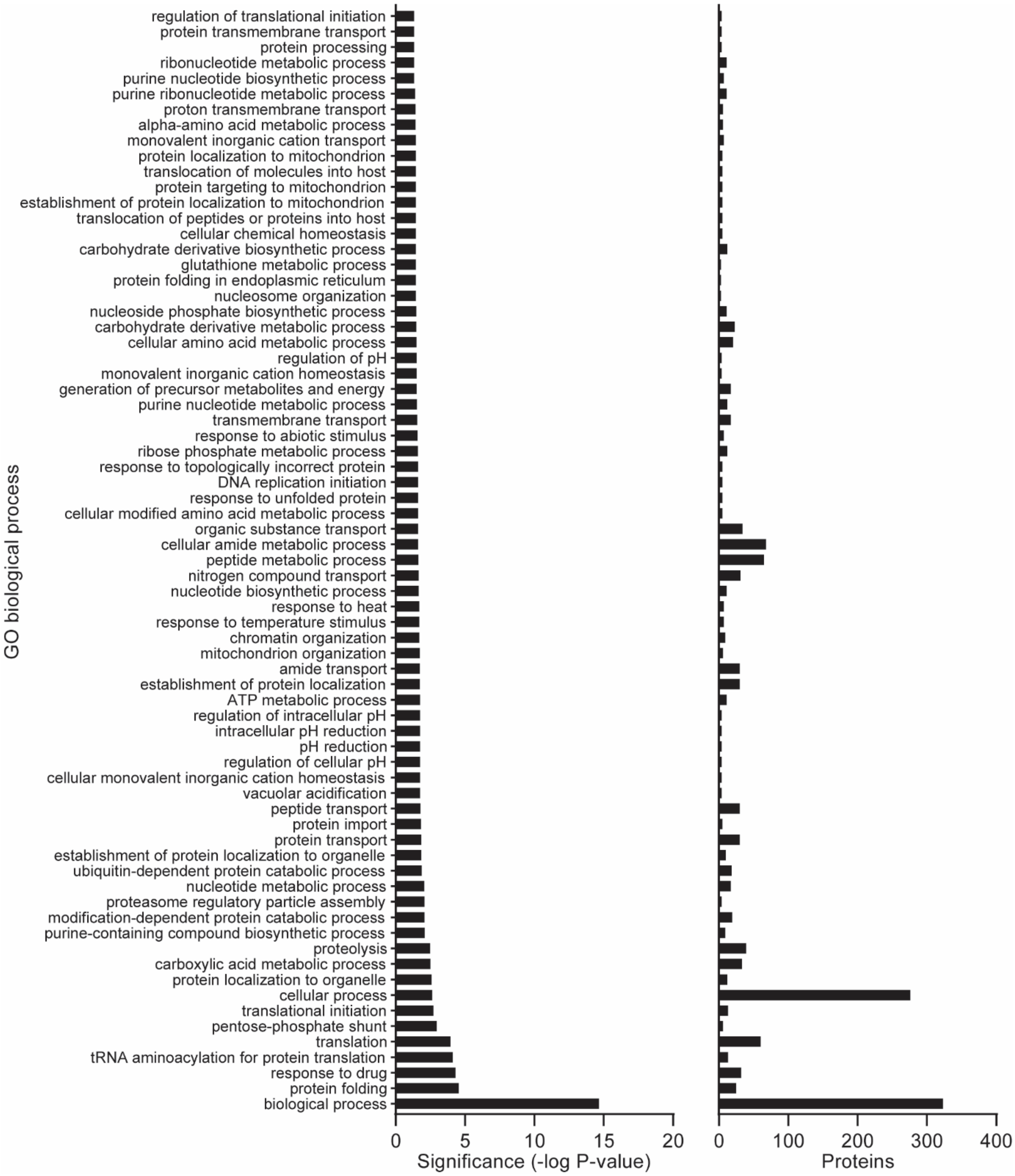


**Figure S3. GO biological process enrichment analysis of control dataset.** Proteins removed by our filtering strategy were subjected to GO analysis for biological process using topGO. Black bars represent the -log P-value for significantly enriched GO biological process terms (left) and the number of proteins identified within each term (right).

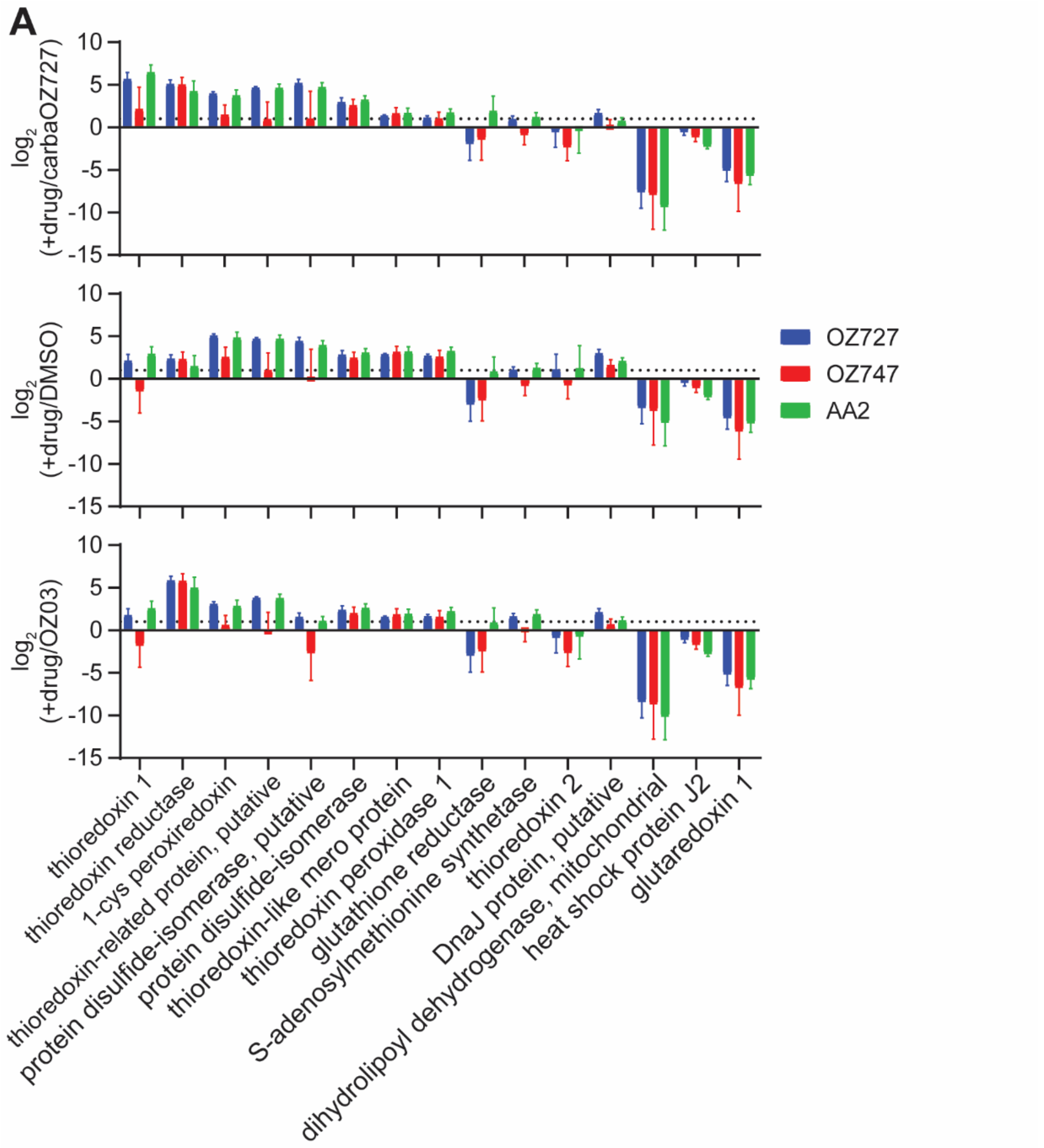
A
log2 (+drug/carbaOZ727)
log2 (+drug/DMSO)
log2 (+drug/OZ03)
10
5
0
-5
-10
-15
OZ727
OZ747
AA2
thioredoxin 1
thioredoxin reductase
1-cys peroxiredoxin
thioredoxin-related protein, putative
protein disulfide-isomerase, putative
protein disulfide-isomerase
thioredoxin-like mero protein
thioredoxin peroxidase 1
glutathione reductase
S-adenosylmethionine synthetase
thioredoxin 2
DnaJ protein, putative
dihydrolipoyl dehydrogenase, mitochondrial
heat shock protein J2
glutaredoxin 1

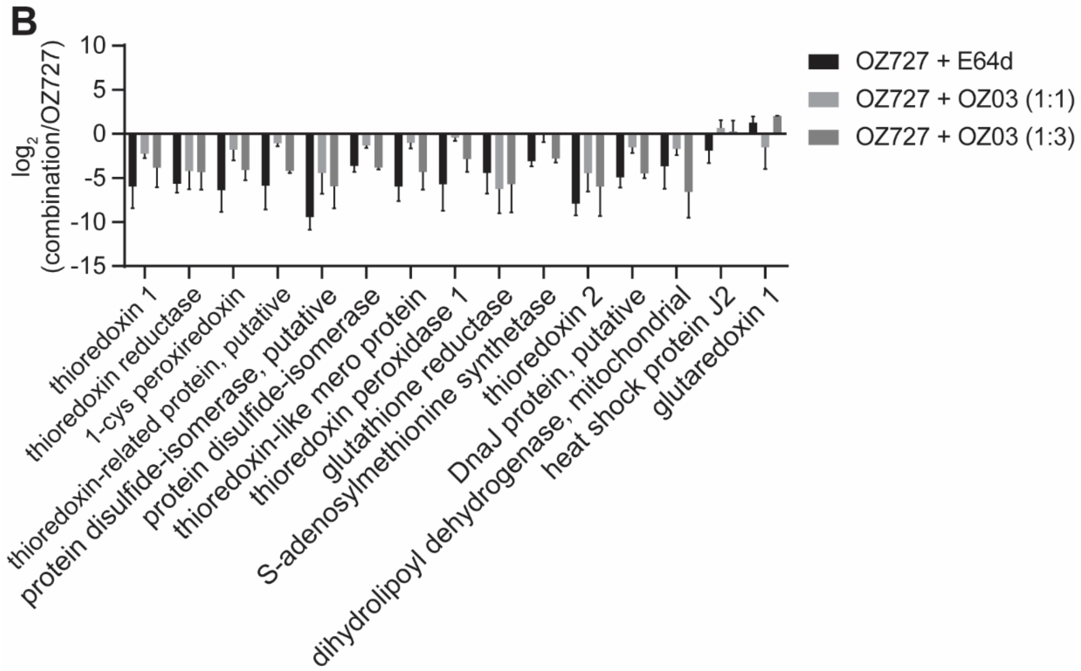
B
log2 (combination/OZ727)
10
5
0
-5
-10
-15
OZ727 + E64d
OZ727 + OZ03 (1:1)
OZ727 + OZ03 (1:3)
thioredoxin 1
thioredoxin reductase
1-cys peroxiredoxin
thioredoxin-related protein, putative
protein disulfide-isomerase, putative
protein disulfide-isomerase
thioredoxin-like mero protein
thioredoxin peroxidase 1
glutathione reductase
S-adenosylmethionine synthetase
thioredoxin 2
DnaJ protein, putative
dihydrolipoyl dehydrogenase, mitochondrial
heat shock protein J2
glutaredoxin 1

**Figure S4. Redox homeostasis proteins (GO:0045454) are targets of peroxide antimalarials.** (A) Enrichment of redox homeostasis proteins (GO:0045454) with OZ727 (blue), OZ747 (red) and AA2 (green) compared to the carbaOZ727 (inactive clickable peroxide), DMSO and OZ03 (active non-clickable parent peroxide) controls. All treatments were for 1 h with 300 nM of the probe. The dashed line represents the two-fold threshold for a peroxide target. (B) Effect of OZ03 co-incubation or E64d pre-treatment on OZ727 labelling of redox homeostasis proteins (GO:0045454). OZ727 activation was blocked by pre-treatment (30 min) of live parasite cultures with 10 µM of the cysteine protease inhibitor, E64d, followed by incubation with OZ727 for an additional 1 h (black). Competition of OZ727 binding to target proteins, was achieved by co-incubation with 300 nM of OZ727 and either an equivalent (300 nM, light grey) or excessive (900 nM, dark grey) concentration of OZ03 for 1 h. The extent of alkylation of redox homeostasis proteins is expressed relative to levels obtained after treatment with 300 nM of OZ727 alone for 1 h. In (A) and (B) data represents the mean ± SEM of at least three biological replicates.

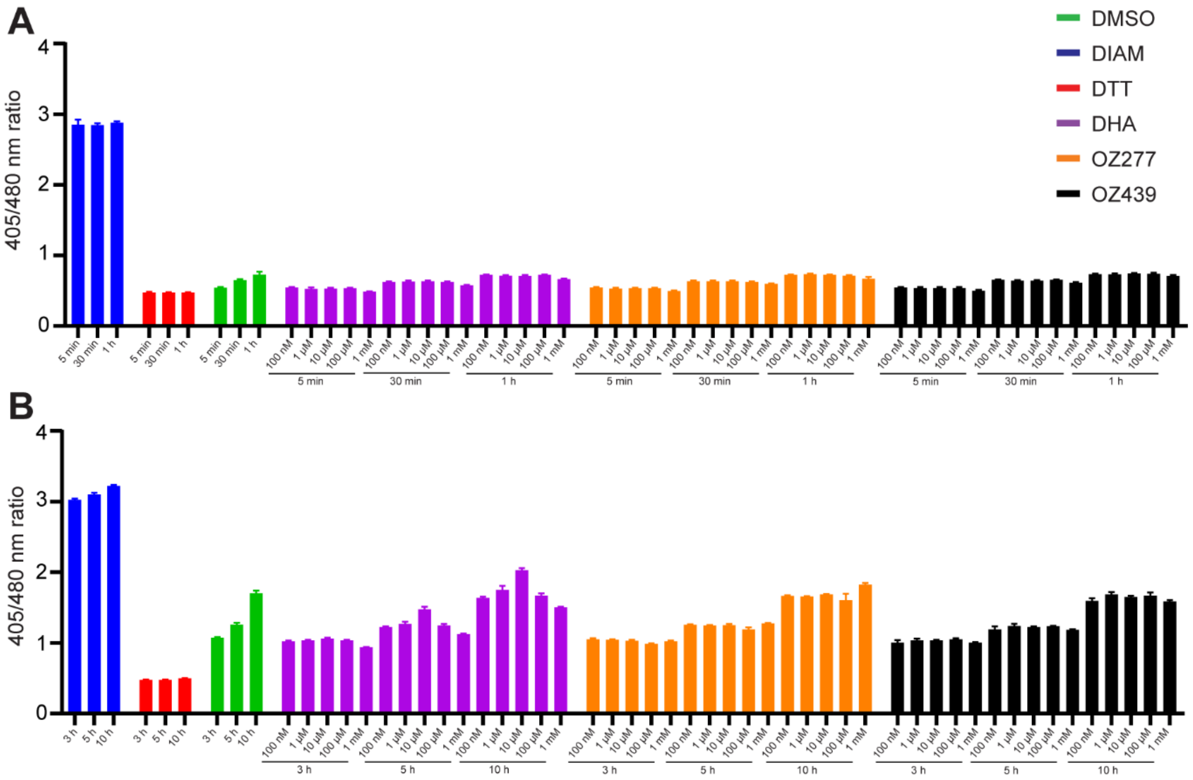


**Figure S5. Effects of peroxide antimalarial on reduced hGrx1-roGFP2.** Reduced recombinant hGrx1-roGFP2 was incubated with peroxides (DHA, OZ277 and OZ439) at increasing concentrations (100 nM, 1 µM, 10 µM, 100 µM and 1 mM). The redox ratio of the recombinant protein (405/480 nm) was measured using a plate reader after 5 min, 30 min, 1 h (A), 3 h, 5 h, and 10 h (B). 10 mM of DTT and 1 mM DIAM was used to fully reduced and oxidized respectively the recombinant protein. Data represents the mean ± SD of two biological replicates, with three technical replicates.

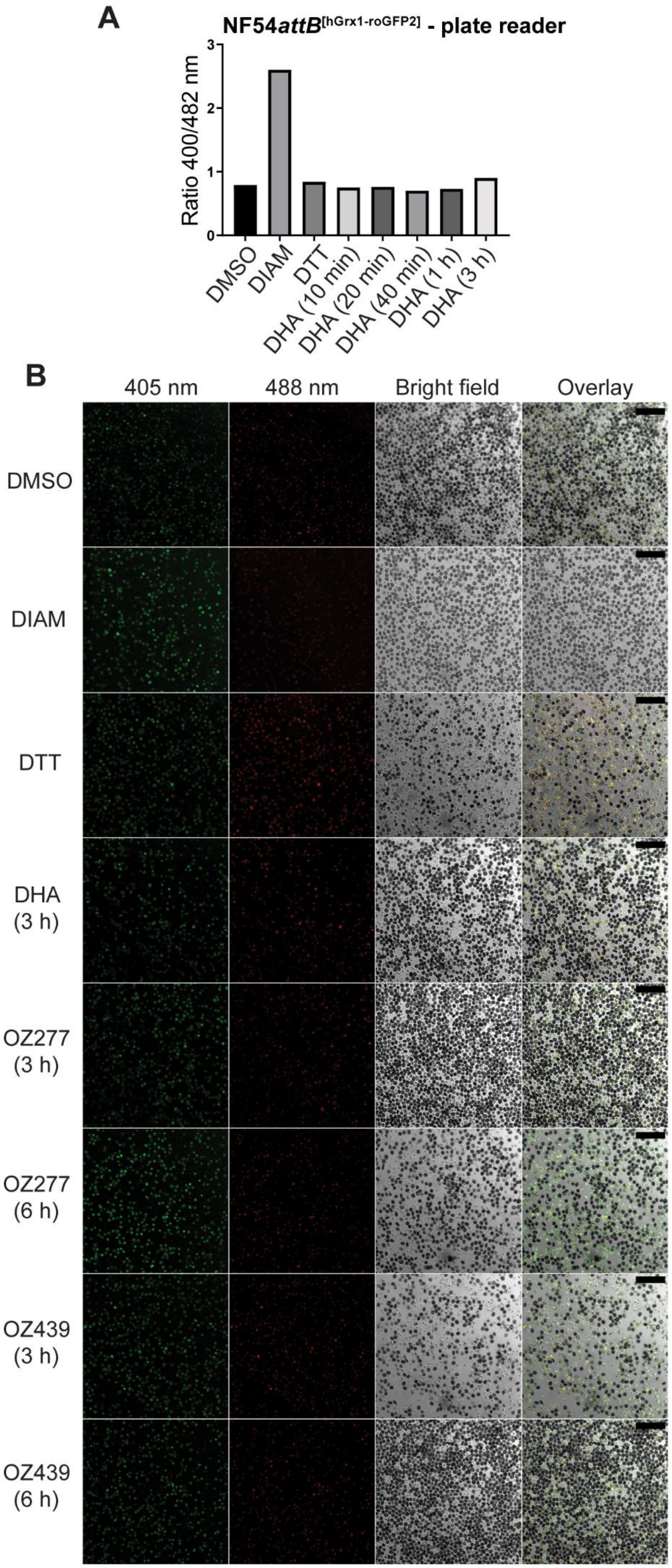
A
NF54attB[hGrx1-roGFP2] - plate reader
Ratio 400/482 nm
0
1
2
3
DMSO
DIAM
DTT
DHA (10 min)
DHA (20 min)
DHA (40 min)
DHA (1 h)
DHA (3 h)
B
405 nm
488 nm
Bright field
Overlay
DMSO
DIAM
DTT
DHA (3 h)
OZ277 (3 h)
OZ277 (6 h)
OZ439 (3 h)
OZ439 (6 h)

**Figure S6. Effect of peroxide antimalarials on the redox ratio of *P. falciparum* NF54*attB*$^{[hGrx1-roGFP2]}$ parasites.** (A) Effects of 100 nM DHA (short incubation - 10, 20, 40 min, 1 and 3 h) on the redox ratio of *P. falciparum* NF54*attB*$^{[hGrx1-roGFP2]}$ parasites. The fluorescence ratio showed that short durations of DHA does not affect the redox sensor within the parasite. Fluorescence was measured using the plate reader and data is from one biological replicate. (B) Parasites were treated with DHA, OZ277 and OZ439 at 300 nM and samples were taken at 3 h post treatment for DHA, OZ277 and OZ439 and at 6 h for OZ277 and OZ439 for the measurement of fluorescence ratio of the redox sensor using CLSM. Fluorescence ratio of the redox sensor increased for parasite treated with OZ277 and OZ439 for 3 and 6 h treatment, while DHA treatment did not change the redox ratio. The black magnification bar represents 50 µm. For both measurements (A and B), DMSO treated parasites at the longest drug duration acted as control, DTT at 10 mM (5 min treatment) was the fully reduced control, while DIAM at 1 mM (5 min treatment) was the fully oxidised control.

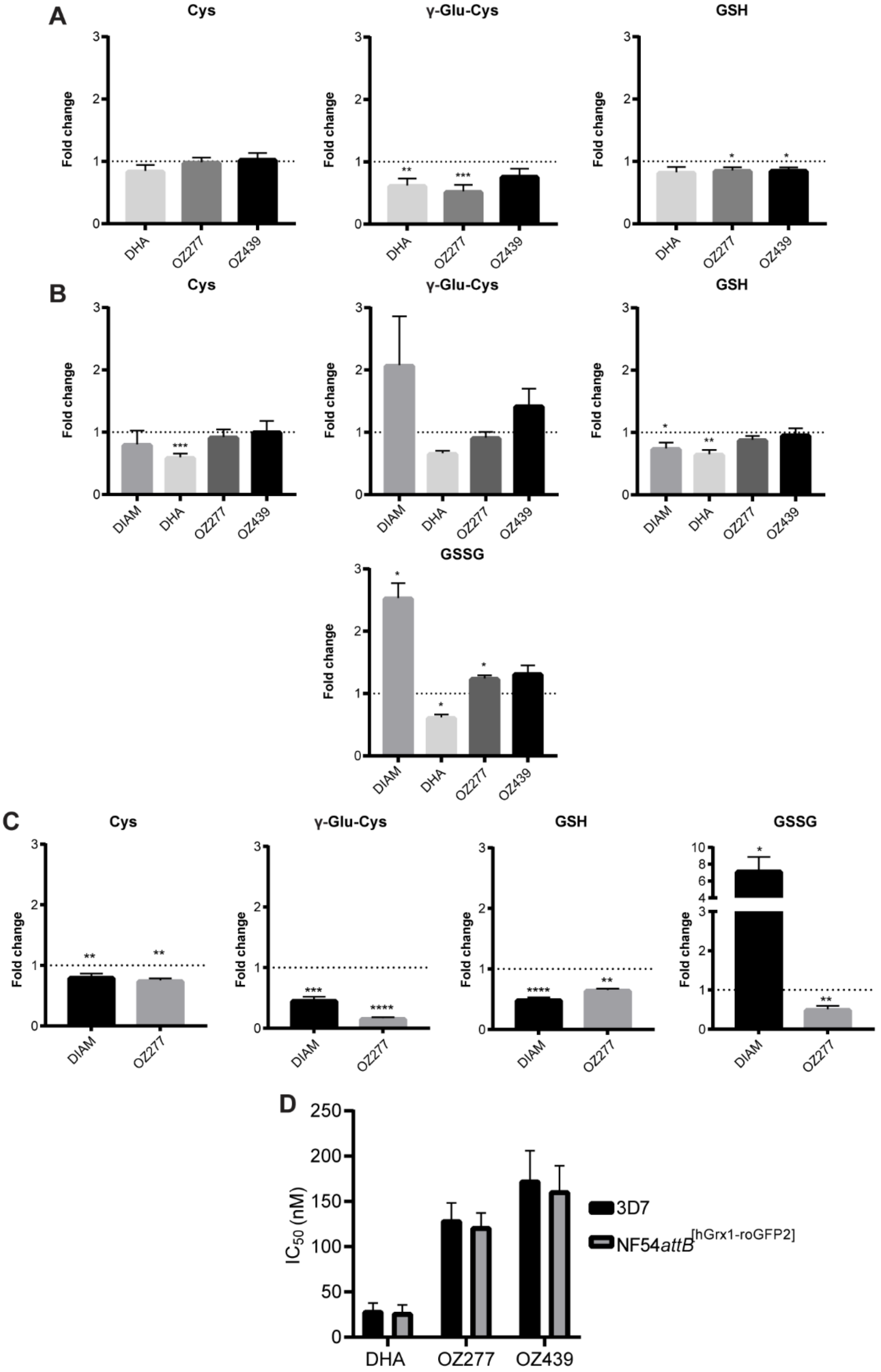


**Figure S7. Effect on thiols (using different sample preparation methods) and activity of peroxide antimalarials in *P. falciparum* 3D7 and NF54*attB*[hGrx1-roGFP2] parasites.** (A, B and C) Determination of relative levels of derivatised cysteine (Cys), γ-glutamyl-cysteine (γ-GluCys), GSH and GSSG compared to DMSO control in *P. falciparum* 3D7 and NF54*attB*[hGrx1-roGFP2] parasites. Treatment with DHA (100 nM, 3 h treatment), OZ277 (300 nM, 3 h treatment), OZ439 (300 nM, 6 h treatment) using method 1 (A) and method 2 (B) in 3D7 parasites, or method 3 (C) in NF54*attB*[hGrx1-roGFP2] parasites.

DTT (10 mM, 5 min treatment) was the fully reduced control, while DIAM (1 mM, 5 min treatment) was the fully oxidised control. Bars represent the relative abundance (mean ± SEM) compared to DMSO control. Total thiol measurements in 3D7 parasites are from two biological replicates (with three technical within each biological) for method 1, and three biologicals replicates (with four technical within each biological) for method 2. Thiol measurement in NF54*attB*$^{[hGrx1\text{-}roGFP2]}$ parasites (method 3) is from one biological replicate (with three technical). P-value was calculated using one-way ANOVA, (*, $p < 0.05$; **, $p < 0.01$, $p < 0.01$; ***, $p < 0.0001$; ****). (D) Activity of peroxide antimalarials on magnet harvested *P. falciparum* 3D7 and NF54*attB*$^{[hGrx1\text{-}roGFP2]}$ parasites. Bar graph showing mean ± SEM $IC_{50}$ values for DHA, OZ277 and OZ439 tested in a 3 h (DHA and OZ277) or 6 h (OZ439) pulse assay with 90% enriched magnet harvested 3D7 and NF54*attB*$^{[hGrx1\text{-}roGFP2]}$ parasites (0.12% Hct). Assays were conducted on three independent occasions in triplicate. There was no significant difference in the measured $IC_{50}$ values for the two parasite lines.
